# Oxidized protein states define metastatic fitness in lung cancer

**DOI:** 10.64898/2026.06.08.730907

**Authors:** Maolin Ge, Magdy Gohar, Consuelo Torrini, Yukako Suzuki, Shan Jiang, Yuchen Huang, Benjamin Weinstein, Janice Wong, Naema Nayyar, Md Yousuf Ali, Emily Sullivan, Stefan Harry, Cagri Cakici, Zhuanglin Dai, Nicolò Vivori, Zhi Qiao, Junbing Zhang, Iphia Zhang, Connie Che, Reilly Stevens, Neha Khandelwal, Jinho Jeong, Yue Xu, Sufian Ibrahim, Haoqian Feng, Philipp Haehnel, Christopher W. Mount, Maria Martinez-Lage, Lecia Sequist, Chendi Li, Sylvia Azer, Quan Lu, Fan-Yan Wei, Mario L. Suvà, Brian B. Liau, Aaron N. Hata, Edward T. Chouchani, A. John Iafrate, Michael S. Lawrence, Doga Gulhan, Priscilla K. Brastianos, Liron Bar-Peled

## Abstract

Reactive oxygen species (ROS) are a pervasive feature of human cancers, yet the protein targets through which ROS-regulated cell states shape tumor biology remains poorly understood. Here, using cysteine chemical proteomics, we define signatures of protein states under distinct cellular ROS environments that capture protein oxidation and conformational changes. Quantifying these signatures in primary lung tumors and brain metastases revealed a surprising enrichment of oxidative states in metastasis. To determine how these states support fitness, we performed genome-wide CRISPR screens and identified the mitochondrial Complex I subunit NDUFA10 as a key oxidation-dependent vulnerability. Oxidation of NDUFA10•Cys253 supports Complex I function through a previously unrecognized nucleotide kinase activity that maintains mitochondrial DNA levels. Enforcing a reduced conformation in NDUFA10 disrupts brain metastatic colonization *in vivo*. These findings establish ROS regulated protein states as a functional layer of tumor fitness, providing a framework for identifying redox-dependent mechanisms that support cancer progression.

## Introduction

Reactive oxygen species (ROS) imbalance is widespread across many cancers, yet their role in tumor biology remains unresolved (*1–5*). ROS have long been viewed as damaging agents that promote genomic instability and tumor initiation (*3, 6–10*), but excessive ROS can also impose a bottleneck to continued tumor growth and metastatic progression (*11*). Indeed, experimental and clinical studies have produced seemingly contradictory conclusions, with ROS reported to both promote and suppress tumor phenotypes depending on context (*12, 13*). Despite this duality, it remains unclear how ROS levels correlate with tumor phenotypes such as aggressiveness or metastatic potential. This question is particularly consequential in lung cancer, where ROS play a key role in cancer development (*3, 14, 15*) and metastasis which is responsible for ∼90% of cancer-related deaths (*13, 16–18*).

A major challenge in understanding ROS biology in tumors is the lack of framework to define and compare cellular redox states in human disease. Most studies infer ROS indirectly through transcriptional activation of antioxidant pathways, measurements of redox metabolites, or genomic scars associated with oxidative stress (*2, 3, 19, 20*). However, these approaches are confounded by pleiotropic functions unrelated to the direct effects of ROS on proteins and incompletely capture how these reactive metabolites impact the functional state of the tumor cell. In contrast to genetic or transcriptional signatures, which robustly define tumor subtypes, vulnerabilities, and prognosis (*21–23*), there is currently no comparable framework for defining ROS-associated cellular states in human cancers. As a result, ROS are typically treated as a nonspecific stress rather than a regulatory signal with defined biological outputs including determinants of cellular state.

Proteins represent a direct molecular interface through which ROS can alter cellular functions (*24–28*). Within the proteome, cysteine residues are uniquely suited to mediate redox regulation due to their chemical reactivity and capacity to undergo reversible oxidation (*24, 26, 29, 30*). Advances in cysteine-focused chemical proteomics have begun to reveal widespread cysteine oxidation in many disease pathologies (*24, 31–36*). Historically, cysteine oxidation has been interpreted as deleterious, largely based on studies showing that oxidation of catalytic cysteines disrupts enzyme activity (*37–42*). However, recent work suggests that oxidation at non-catalytic or allosteric cysteines can not only alter catalytic activity, but also protein conformation and interaction state (*41, 43–48*). Thus, coordinated changes in cysteine accessibility across the proteome which we refer to as protein states, may encode information about the underlying redox environment of a cell in a manner conceptually analogous to how coordinated transcriptional programs define gene-expression signatures (*22*). However, whether oxidation more specifically affects protein conformation, functional regulation, or disease phenotypes in human tumors remains unknown.

Here, we sought to determine the oxidized and reduced protein states that lung cancers inhabit, track how these states change during cancer progression, and identify ROS-dependent mechanisms that support tumor fitness. To answer these questions, we developed an experimental framework that systematically perturbs intracellular ROS levels across a panel of 55 non-small cell lung cancer (NSCLC) cell lines and measures proteome-wide cysteine accessibility using chemical proteomics. By integrating changes in cysteine oxidation and protein conformation, this approach allowed us to derive defined oxidized and reduced protein-state signatures.

Quantifying these signatures against cysteine accessibility profiles of 31 primary lung tumors with matched adjacent normal tissue and 19 brain metastases unexpectedly revealed that the majority of metastatic lesions are enriched for oxidized protein states. This is in contrast to the prevailing notion that high levels of oxidation are disabling to tumor growth and instead suggest that oxidized cellular environments support functions essential for metastatic progression. To identify such functions, we combined ROS-state profiling with genome-wide CRISPR screens and uncovered a dependency on oxidation-sensitive mitochondrial Complex I function. Mechanistically, we find that higher-order oxidation of NDUFA10•Cys253 to a sulfinic acid supports Complex I function and a previously unrecognized nucleotide kinase activity that is necessary for dGTP production and mitochondrial DNA (mtDNA) maintenance. Enforcing a constitutively reduced conformation at this residue in lung cancer models blocks brain metastatic colonization and suppresses growth *in vivo*.

Collectively, these findings establish ROS-associated protein states as a functional signature of tumor cell regulation in lung cancers, revealing that metastases to the brain are highly oxidized. Importantly, we identify one mechanism by which oxidation-dependent protein conformations support metastatic fitness, providing a scaffold for systematically identifying ROS-dependent regulatory mechanisms across tumor progression and other disease contexts.

## Results

### Defining ROS-associated protein conformational states in lung cancer

ROS levels across stages of lung cancer progression have been inferred through transcriptional programs such as NRF2 and NF-κB activation, or through genomic scars induced by oxidative stress (*17, 49–51*). However, how ROS impacts global protein states across different stages of lung cancer remains unknown. Because proteins are direct effectors of cellular function, we reasoned that defining tumor states at the level of protein oxidation and conformation, could provide a mechanistic framework linking ROS environments to cancer phenotypes.

To directly address this question, we profiled proteome-wide cysteine accessibility using chemical proteomics in human lung cancer specimens. Cysteine accessibility is shaped both by direct oxidation (e.g., sulfinylation) (*31, 35, 52*) and changes in protein conformation that alter solvent exposure (*53*). Coordinated cysteine accessibility changes across the proteome can be interpreted as *signatures* that reflect the combined effects of oxidation and structural remodeling to a specific cellular environment; we refer to these as protein states. We therefore sought to define ROS-associated protein states from a structural perspective using cysteine accessibility signatures analogous to gene-expression signatures used to classify transcriptional programs in tumors (*21, 54*)(Fig. 1A).

**Fig. 1:**
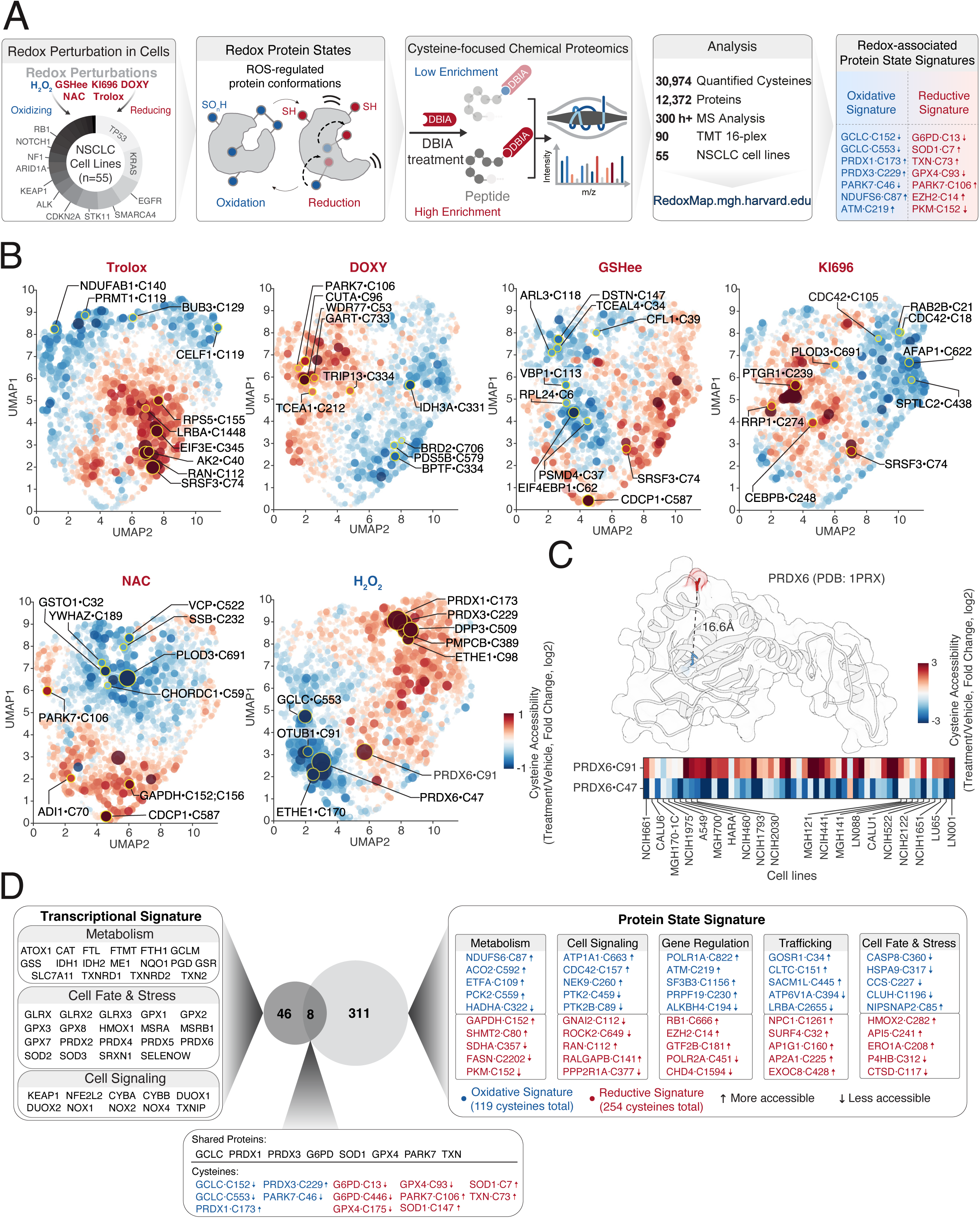
**Defining redox-associated protein states in lung cancer**. (**A**) Chemical proteomic workflow to assess protein states following changes in intracellular ROS levels across non-small cell lung cancer (NSCLC) models. 55 NSCLC cell lines were treated with the indicated redox-state perturbations: 1 µM KI696 (NRF2 activator)(*106*), or 5 µM DOXY (mitochondrial translation inhibitor) for 48 hrs; 10 mM NAC, 10 mM GSHee (thiol-buffering agents), 250 µM Trolox (radical-trapping antioxidant) for 1 hr; or 200 µM H_2_O_2_ for 30 min. Cells were harvested, lysed, and treated with a cysteine-specific desthiobiotin iodoacetamide (DBIA) probe, and relative levels of streptavidin-enriched cysteines were determined by quantitative TMT-based mass spectrometry (see also Methods). (**B**) Cysteine accessibility increases and decreases upon changes in ROS levels. UMAP plots depicting changes in cysteine accessibility upon different ROS perturbations (see also Table S1). (**C**) H_2_O_2_ alters the conformation of PRDX6. Top, structural representation of PRDX6 (PDB:1PRX) highlighting the location of active-site Cys47 and peripheral Cys91. Bottom, heatmap displaying opposing accessibility changes for each cysteine in PRDX6 following H_2_O_2_ treatment across cell lines profiled. (**D**) Defining redox-associated protein state signatures. Overlap of transcriptional signatures for ROS (GSEA = Reactive oxygen species pathway) (*22*) with protein-state signatures developed herein. Protein-state signatures are composites of cysteine accessibility changes following perturbations that increase (blue) or decrease (red) intracellular ROS (see also Table S4, Methods).

To explicitly associate these signatures with changes in cellular redox environment rather than unrelated cellular states, we developed a reference atlas of cysteine accessibility changes under defined ROS perturbations. We modulated cellular ROS levels across a panel of 55 NSCLC cell lines representing diverse oncogenic genotypes (Fig. 1A, fig. S1A-C). We quantified cysteine accessibility by treating lysates with the cysteine-reactive electrophile desthiobiotin iodoacetamide (DBIA) (*48, 55*) followed by isoTMT-based chemical proteomics (*55–58*) (Fig. 1A, Table S1). To generate an oxidized cellular environment, cells were treated with H_2_O_2_. To induce more reduced states, we modulated antioxidant programs by: (1) activating NRF2, a master regulator of antioxidant responses (*51, 59, 60*), and (2) inhibiting mitochondrial translation, a central mechanism used by cells to regulate mitochondrial ROS (*48, 61*) (fig. S1A). In parallel, we treated cells with small molecule reducing agents, including the thiol-buffering agents N-acetylcysteine (NAC) and glutathione ethyl ester (GSHee), and the radical-scavenging antioxidant Trolox (*62–64*) (fig. S1A), which collectively decreased intracellular ROS as measured with the H_2_DCFDA probe (fig. S1D). Because some perturbations extended beyond 24 hours, we additionally quantified protein expression changes across 19 NSCLC cell lines to identify proteins whose abundance changed following treatment. These proteins were excluded from subsequent chemical proteomic analyses, allowing cysteine accessibility changes to be attributed specifically to redox perturbations as opposed to expression differences (fig. S1E, Table S2, and Methods). At the global level, each perturbation impacted distinct regions of the cysteinome (Fig. 1B, fig. S1F), consistent with their differing chemical and biological mechanisms. Cysteine accessibility changes evoked by H_2_O_2_ showed an opposing pattern to those following NRF2 activation, whereas NAC and GSHee largely clustered together, consistent with their shared thiol-buffering properties (Fig. 1B, fig. S1F). Importantly, each perturbation induced both increases and decreases in accessibility across the cohort of cells profiled (Fig. 1B, fig. S1C, G), consistent with redox-linked conformational remodeling of protein states rather than purely an unidirectional change in cysteine oxidation state. For example, H_2_O_2_ treatment decreased accessibility of the catalytic cysteine (C47) in PRDX6, a well-established target of H_2_O_2_ oxidation (*65, 66*) (Fig. 1B, C). In contrast, accessibility at a distal cysteine in PRDX6 (C91), located ∼16.6 Å from the active site, was simultaneously increased by H_2_O_2_, consistent with redox-induced conformational rearrangement (Fig. 1C). Likewise, perturbations that lowered intracellular ROS such as GSHee, increased cysteine accessibility across the proteome as expected, but also decreased cysteine accessibility at distinct subsets of cysteines, further supporting the premise that this approach captures proteome-wide structural remodeling in addition to direct cysteine oxidation (Fig 1B, fig. S1C, G). We make this dataset available via a searchable online interface, to enable studies of protein oxidation and redox-associated conformational regulation.

To understand how distinct redox environments reshape the proteome, we used cysteine set enrichment analysis (CSEA), a computational pipeline to determine enrichment of cysteine accessibility changes across pathways, organelles and protein classes (Table S3) (*53*). To assess the overall impact of protein-state changes across these gene sets, cysteines were ranked by the absolute accessibility change weighted by the negative log10-transformed p-value, and enrichment analysis was performed on the resulting scores. CSEA revealed that different redox perturbations impacted cysteines at distinct cellular locations, with H_2_O_2_ and GSH prominently affecting mitochondrial cysteines (fig. S1H). Increasing ROS levels revealed accessibility changes in nuclear proteins, suggesting organelle-specific sensitivity to redox environment (fig. S1H). CSEA analysis further identified specific protein classes and pathways impacted by each perturbation, with accessibility changes induced by H₂O₂ treatment were enriched in oxidoreductases, whereas NRF2 activation enriched in oxidoreductases and the corresponding ferroptosis pathway (fig. S1I, Table S3). Together, these results reveal reproducible and conserved accessibility patterns that collectively define distinct oxidized and reduced protein states.

To define oxidative and reductive signatures, we selected cysteines with accessibility changes ≥ 0.5 or ≤ -0.5 and p-value ≤ 1 × 10^-6^ in at least one of the conditions. Collectively, 381 cysteines spanning 319 proteins contribute to these signatures spanning different cellular pathways and functions (Fig. 1D, Table S4, and Methods). As expected, known-regulated cysteines were recovered, including PKM•C423/C424 and CCS•C227, which exhibited decreased accessibility under H_2_O_2_ treatment, and GAPDH•C152, which exhibited increased accessibility under reducing perturbations (*67–69*). Interestingly, these signatures were mostly non-overlapping (Table S4), enforcing the premise that they capture unique cellular environments, rather than a continuum of oxidative stress. Comparisons with traditional transcriptional signatures revealed partial overlap between cysteines and their corresponding genes (Fig. 1D), highlighting distinct layers of redox cell states captured by each modality: transcriptomics reports the activity of transcription factors, whereas chemical proteomics captures protein states.

### Oxidized protein states are enriched in brain metastases

We next asked whether primary lung tumors and metastatic lesions occupy distinct oxidized or reduced states by analyzing a cohort of 31 tumor–normal paired lung cancer specimens together with 19 brain metastases collected from patients treated at Mass General Hospital (Fig. 2A, fig. S2A, Table S5). These tumors represented common NSCLC driver genotypes (TP53 40%, EGFR 32%, KRAS 26%, ALK 6% (Fig. 2B). Specimens were cryo-milled to minimize artifactual oxidation and subjected to cysteine accessibility profiling using the workflow described above (Fig. 2A). To account for differences in protein abundance across samples, accessibility changes were normalized to total proteome quantification across all samples (Methods). To account for tumor purity and contamination from normal tissues, we developed tissue-specific protein expression signatures using stroma, immune, and brain protein sets and excluded cysteines whose accessibility correlated with these signatures (fig. S2B, C, Methods). Confirming the validity of normal tissue protein expression signatures, we found brain signature to be elevated in brain metastases, and stroma signature to be higher in primary tumors, reflecting expected differences in the associated tumor microenvironment.

**Fig. 2:**
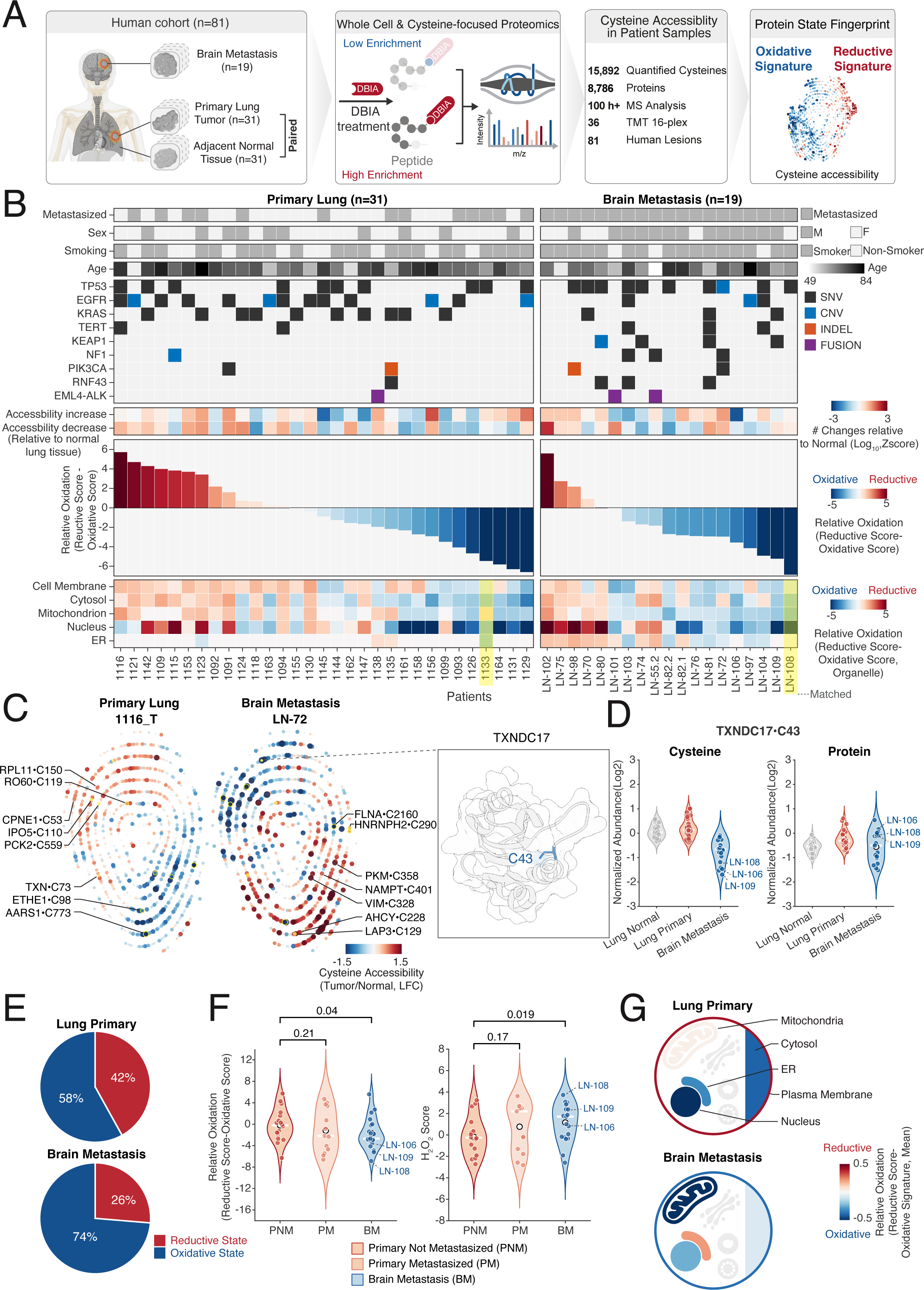
Lung Cancer Brain Metastases are enriched for oxidized protein states. (**A**) Chemical proteomic workflow to assess cysteine accessibility and protein state signatures in primary human lung tumors. (**B**) Relative oxidation in primary lung lesion and brain metastases. From top to bottom: heatmap listing clinical features of patient cohort analyzed in this study (see also Table S5); Heatmap depicts number of cysteine accessibility changes relative to normal lung tissue; Waterfall plot indicates relative oxidation in each specimen by subtracting the reductive signature score from the oxidative signature score (see also fig. S3A-B, Methods); Compartment specific heatmap displaying NES for oxidized (blue) or reductive (red) signatures for each organelle. (**C**) Primary lung and brain metastases have cysteine accessibility differences relative to matched normal tissue. Left, fingerprint plot displaying cysteine accessibility changes between primary lung tumor (1116_T) and brain metastasis (LN-72) relative to normal lung tissue. Each dot represents a cysteine. Right, structure of TXNDC17 (model: AF-Q9BRA2-F1), highlighting the location of Cys 43. (**D**) Violin plot depicting changes in cysteine accessibility for TXNDC17•C43 and corresponding protein levels in patient samples. (**E**) Pie charts depicting oxidative and reductive states in specimens profiled. (**F**) Metastatic lung tumors and brain metastases are more oxidative than non-metastatic primary lung tumors. Left, violin plot depicting relative protein state oxidation across cohort. Right, enrichment scores for H_2_O_2_ signature in each group of specimens profiled in this study. (**G**) Mitochondria display oxidized protein states in brain metastases. Cellular heatmap depicting relative protein states at each organelle. Statistical significance was determined by one-tailed Wilcoxon rank-sum test.

Cysteine accessibility profiling revealed robust and reproducible differences between normal lung tissue, primary tumors, and brain metastases that were not explained by protein abundance alone (Fig. 2B, fig. S2D). To examine how cysteine accessibility reflects tumor protein states at the patient level, we generated cysteine accessibility fingerprints for representative primary lung tumors and brain metastases (Fig. 2C), providing residue-level insight into tumor-driven remodeling of the proteome. Primary tumors and brain metastases exhibited distinct accessibility landscapes, with individual cysteines showing stage-specific alterations relative to normal lung tissue. For example, TXNDC17•C43 exhibited reduced cysteine accessibility in brain metastases compared with primary tumors, whereas total TXNDC17 protein abundance was largely unchanged across disease states (Fig. 2D). Thus, cysteine accessibility profiling identified large-scale conformational remodeling in tumors compared normal tissue, as well as, between different stages of cancer.

We applied our experimentally derived signatures across primary tumors and metastatic lesions to determine whether metastatic progression is associated with distinct ROS-associated states. Cysteine accessibility changes in tumors were determined relative to that in normal lung tissue and quantified in single-sample cysteine enrichment analysis (ssCSEA) using the experimental cysteine sets (Methods). Relative oxidation for each tumor was calculated as the difference between composite reductive and oxidative enrichment scores, with positive scores indicating more reductive states and negative scores indicating more oxidative states (fig. S3A). Reassuringly, we observed a robust relative oxidation signal driven by both enrichment of oxidative signatures and reductive signatures across samples. Known redox-sensitive cysteines, such as TXN•C73, exhibited increased accessibility in more reductive states, supporting the biological validity of the inferred redox scores (fig. S3B). We observed global shifts in relative oxidation levels, with increasingly oxidized tumors displaying coordinated remodeling across cysteine sites (Fig. 2B, fig. S3B-C). Moreover, tumors with similar oxidation states exhibited highly concordant proteome-wide cysteine accessibility patterns, highlighting the pervasive influence of redox status on protein-state organization (fig. S3C). Strikingly, the majority of metastatic lesions were in oxidized protein states (n=15/19), whereas oxidized and reduced proteins states were more equally distributed across primary tumors (Fig. 2B, E, fig. S3B). To determine if relative oxidation tracked with clinical features, primary tumors were further segregated into those that went onto metastasize and those that did not. Comparing relative oxidation between the subsets indicated that primary tumors that metastasized were more similar in their oxidation state to metastases. In contrast, primary tumors that did not go on to metastasize were significantly less oxidized than metastases (p=0.045) (Fig. 2F). Consistent with overall redox status of the tumor cohorts, the H_2_O_2_ specific signature was significantly enriched in metastatic lesions compared to primary tumors that did not metastasize (p=0.019) (Fig. 2F). These results indicate that metastatic progression is associated with increasingly oxidized proteomic environments.

The established cellular localization of proteins comprising our redox signatures allowed us to ask how ROS-associated protein states change between cellular compartments. While the nucleus was largely oxidized across specimens in both tumor types, mitochondria were, on average, more enriched for oxidized protein states in metastases compared to primary tumors (Fig. 2G). In one patient with matched primary and metastatic lesions, mitochondria shifted from reductive proteins states to oxidative, despite both tumors maintaining an overall oxidizing environment (Fig. 2B). Comparison of mitochondrial protein states revealed that metastases were significantly more oxidized than primary tumors that did not metastasize (p=0.025) (fig. S2E). Together, these data establish four central observations: primary tumors differ substantially from adjacent normal lung tissue at the level of cysteine accessibility, brain metastases occupy distinct proteomic states relative to primary tumors, experimentally defined oxidized protein states are preferentially quantified in metastatic lesions and the mitochondria is enriched for oxidized protein states in metastasis. Collectively, these data suggest that oxidized protein states may not simply be tolerated during tumor progression, but may instead be selected as functional states that support metastatic fitness.

### Disruption of mitochondrial Complex I mediates sensitivity to a reduced cell state

Our analysis of patient lung tumors revealed that brain metastases are enriched for oxidized protein states, suggesting that the corresponding environment is not merely a byproduct of oxidative stress, but may instead be selectively maintained to support tumor fitness. We therefore hypothesized that metastatic cells may become dependent on proteins whose function requires oxidation-dependent conformational states. To identify pathways and genes required for fitness under oxidized conditions, we sought genes whose loss mediates cellular sensitivity when ROS levels are lowered.

To address this question, we performed functional genomic CRISPR screens under conditions that shift cells toward reduced protein states (Fig. 3A). Among NSCLC models examined, CALU6 cells exhibited one of the strongest changes in cysteine accessibility following treatments that lower ROS and exhibited pronounced sensitivity to these pertubations in proliferation assays (fig. S4A, B), making it a tractable system for genetic interaction mapping. CALU6 cells transduced with a genome-wide sgRNA library were cultured for 11 population doublings under vehicle or in the presence of five distinct ROS decreasing conditions. Distinct genes and pathways mediating sensitivity and resistance were identified for each treatment (Fig. 3B, fig. S4C, Table S6). For example, loss of the cytosolic peroxidase PRDX1 (*70, 71*) conferred resistance to multiple treatments which decrease ROS (fig. S4C), consistent with its role in peroxide detoxification and the importance of maintaining oxidized environments. Strikingly, the electron transport chain (ETC) emerged as a dominant determinant of sensitivity to lower ROS, with Complex I genes standing out across multiple treatments (Fig. 3B-D, fig. S4C).

**Fig. 3:**
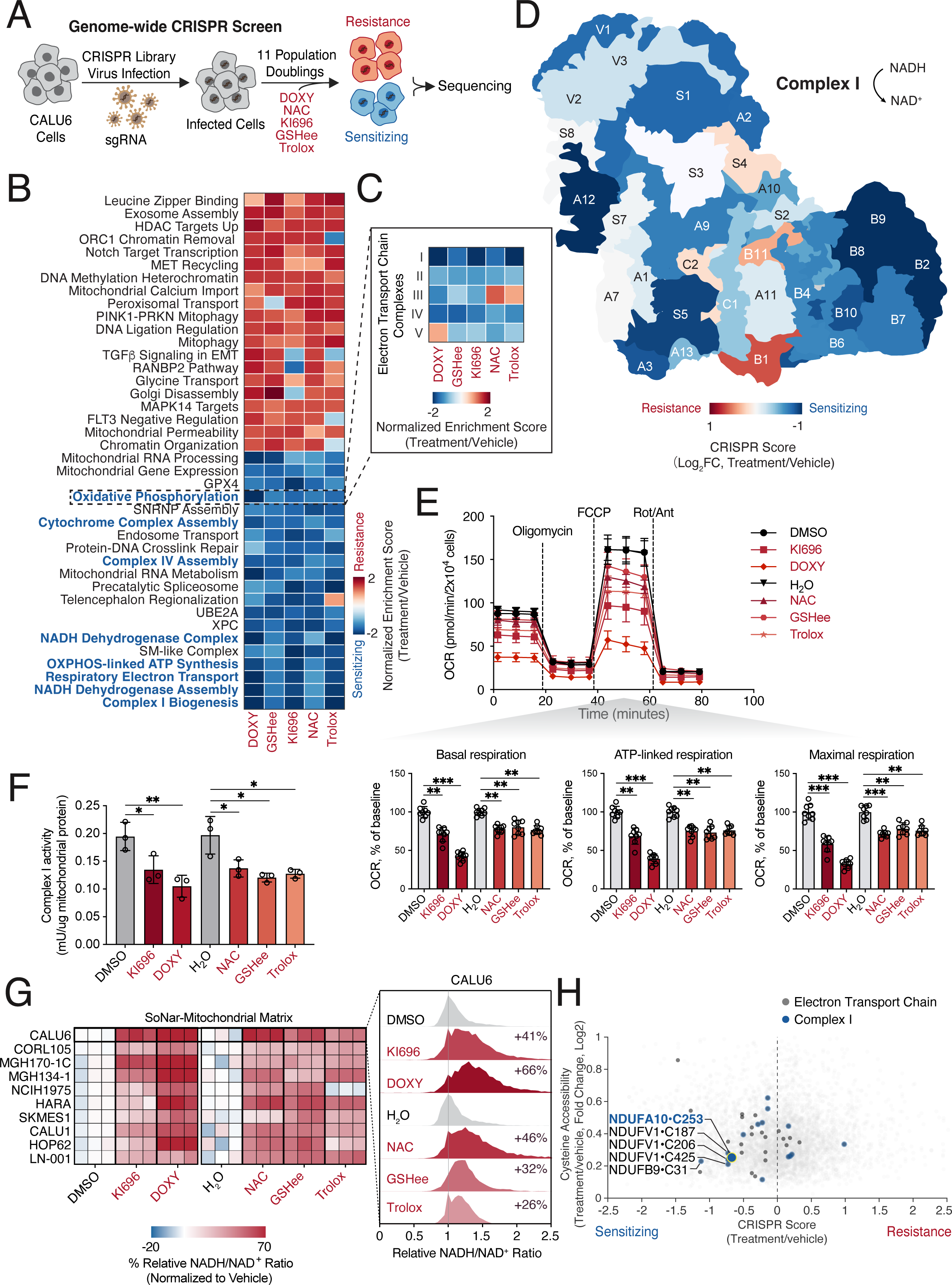
Loss of mitochondrial Complex I sensitizes NSCLCs to reduced cell states. (**A**) Schematic of functional genomics analysis to identify genes and pathways that mediate sensitivity to a reduced cellular state. CALU6 cells were transduced with a genome-wide sgRNA library and cultured for 11 doublings in the presence of the indicated compounds that promote a reduced cell state: 1 µM KI696, 2.5 µM DOXY, 5 mM NAC, 5 mM GSHee, or 125 µM Trolox. sgRNA counts were determined by sequencing and used to calculate CRISPR scores (see also Table S6 and Methods). Negative CRISPR scores (in blue) indicate genes whose loss sensitizes cells to the indicated perturbation, and positive scores (in red) indicate genes whose loss provides resistance. (**B**) Loss of oxidative phosphorylation (OXPHOS) mediates sensitivity across reduced cell states. Heatmap depicts normalized enrichment scores (NES) for cellular pathways (*22*). (**C**) Mitochondrial Complex I loss drives sensitivity to reduced cell states. Heatmap depicts normalized enrichment scores for each electron transport chain (ETC) complex for corresponding reducing conditions. (**D**) Median CRISPR scores of ETC Complex I subunits mapped onto the structure of mammalian Complex I (PDB: 5XTD) (*107*). (**E**) A reduced cell state impairs mitochondrial respiration. CALU6 cells were treated with 1 µM KI696 or 5 µM DOXY for 48 hrs, or 10 mM NAC, 10 mM GSHee, and 250 µM Trolox for 1 hr, and oxygen consumption rate (OCR) was measured by Seahorse analysis. Representative OCR traces are shown, with quantification of basal respiration, ATP-linked respiration, and maximal respiration displayed below. (**F**) A reduced cell state suppresses mitochondrial Complex I activity. CALU6 cells were treated as described in (**E**) and Complex I activity was measured following the addition of NADH and ubiquinone_1_ to isolated mitochondria membranes. Bars indicate Complex I activity normalized to total mitochondrial protein. (**G**) Decreasing ROS levels with reduced-state perturbations increases the mitochondrial NADH/NAD^+^ ratio. Cells expressing SoNar targeted to the mitochondrial matrix were treated as described in (**E**), and mitochondrial NADH/NAD^+^ was measured by FACS. Left, heatmap showing percent change in the mitochondrial NADH/NAD^+^ ratio across NSCLC cell lines relative to vehicle. Right, representative FACS plots from CALU6 cells expressing SoNar. The NADH/NAD^+^ ratio was determined by comparing the emission intensity of 405 nm (NADH binding) vs. 488 nm (NAD^+^ binding) for SoNar. (**H**) Integration of cysteine chemical proteomics and functional genomics prioritizes NDUFA10•C253 as a top target. Scatter plot comparing max magnitude of cysteine accessibility change across reduced-state perturbations with corresponding CRISPR score (see also Methods). Statistical significance was determined by Student’s *t* test (two-tailed, unpaired). * *p* < 0.05; ** *p* < 0.01; *** *p* < 0.001.

Because patient metastases exhibited oxidized protein state signatures, together with the convergence of our CRISPR screens on Complex I, led us to ask whether lowering ROS directly impairs mitochondrial respiration. We treated CALU6 and MGH134-1 cells with multiple agents that decrease ROS and measured mitochondrial respiration using Seahorse analysis. These treatments reduced basal respiration, ATP-linked respiration, and maximal respiration (Fig. 3E, fig. S5A). These effects were accompanied by decreased Complex I activity (Fig. 3F, fig. S5B) and by increased NADH/NAD^+^ ratios measured using the genetically encoded SoNar reporter (*72*) targeted to either the mitochondria matrix or cytosol across multiple NSCLC cell lines (Fig. 3G, fig. S5C). Excessive ROS was also detrimental to mitochondrial respiration, as high-dose H_2_O_2_ similarly impaired oxygen consumption (fig. S5D, E), consistent with prior observations (*73, 74*). Together, these findings suggest that Complex I activity requires a balanced redox environment to function, and that oxidized states appear to support mitochondrial Complex I activity, thereby creating a selective dependency on oxidation-sensitive mitochondrial functions.

### NDUFA10 oxidation induces a conformational change and supports Complex I function

Having identified Complex I as a key dependency of cancer cells under reducing environments, we next sought to determine the molecular mechanism linking ROS states to Complex I function. We reasoned that proteins requiring oxidation-dependent conformations for activity should satisfy two criteria: 1) they should undergo ROS-associated cysteine accessibility changes in our proteomic atlas, and 2) their function should become limiting when cells are forced into reduced states.

To identify such candidates, we integrated our chemical proteomic analysis of ROS cell states with CRISPR loss-of-function screening to nominate high-priority cysteine targets that may need oxidation for function (Fig. 3H, Methods). Focusing on the ETC given the results of our CRISPR screen, this analysis highlighted 17 candidates, with the Complex I cysteine NDUFA10•C253 emerging as one of the top-ranked hits (Fig. 3H and 4A). NDUFA10 is required for Complex I assembly (*75*), and consistent with this role, depletion of NDUFA10 disrupted mitochondrial respiration and Complex I activity, and increased NADH levels, a phenotype rescued by expression of an sgRNA-resistant NDUFA10 PAM mutant (Fig. 4B-E, fig. S6A-E). To determine whether the redox state of Cys253 directly regulates NDUFA10 function, we generated Cys–>Ser and Cys–>Asp mutants of reduced and higher-order oxidized (sulfinic acid-like) cysteines, respectively (*61, 76*) (Fig. 4B). While Asp preserves the negative charge associated with cysteine sulfinic acid, it differs in its geometry and coordination chemistry. Therefore, we interpret our findings with this mutant as a mimetic for this type of ROS post-translational modification. NDUFA10•C253S failed to rescue the NADH/NAD^+^ imbalance and Complex I defects caused by NDUFA10 loss (Fig. 4C-E, fig. S6D, E). In contrast, the NDUFA10•C253D mutant restored the NADH/NAD^+^ ratio and Complex I activity to near wild-type levels (Fig. 4C-E, fig. S6D, E). Moreover, cells expressing the reduced mutant markedly suppressed respiration relative to WT and the oxidized mimetic (Fig. 4C). Defective Complex I activity caused by reduction of NDUFA10•C253 was similarly reproduced across multiple cell lines using traditional Complex I activity assays (Fig. 4D). Consistent with a functional requirement for NDUFA10 oxidation, expression of the NDUFA10•C253D mutant partially reversed NADH reductive stress (*77*) and mitigated partial Complex I inhibition following a decrease in ROS levels and NRF2 activation (Fig. 4F, fig. S6F, G).

**Fig. 4:**
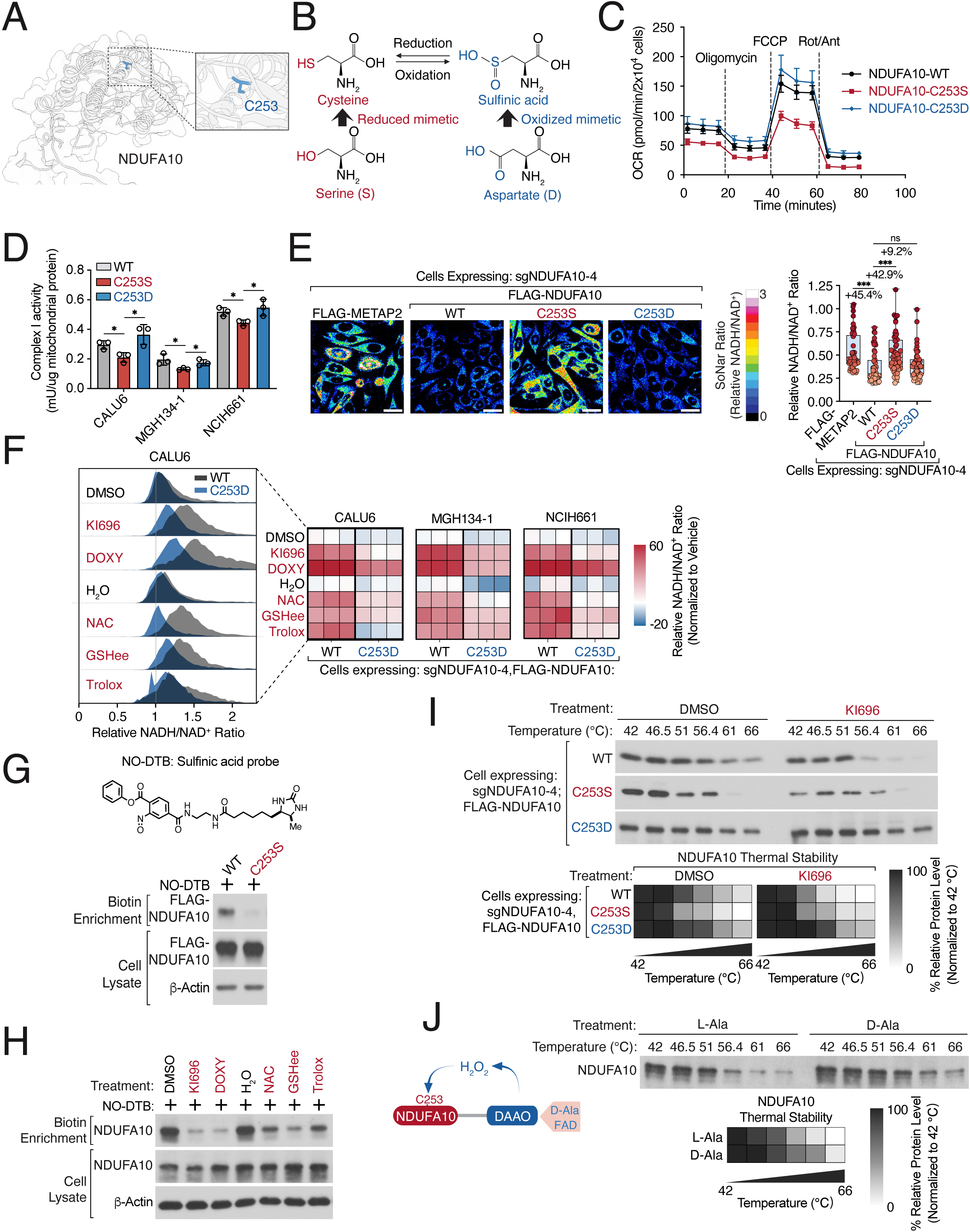
Oxidation of NDUFA10 induces a conformational change and promotes Complex I function. (**A**) Structure of NDUFA10 (model: AF-O95299-F1) with Cys253 highlighted. (**B**) Amino acid redox mimetics, that complement cysteine in its reduced (Ser) and oxidized (Asp) states. (**C**) Differential mitochondrial activity in cells expressing NDUFA10-redox mimetics. Seahorse analysis of cells depleted of NDUFA10 and co-expressing the indicated PAM-mutant NDUFA10 in its wild-type (WT)– or redox variant state. (**D**) Oxidation of NDUFA10 enhances Complex I activity. Complex I activity was measured in the indicated cell lines depleted of NDUFA10 and co-expressing the indicated PAM-mutant NDUFA10 variants as described in (**B**). (E) The sulfinic acid mimetic of NDUFA10 is sufficient to support NADH oxidation. Left, representative immunofluorescence images of CALU6 cells depleted of NDUFA10 and co-expressing the indicated PAM-mutant NDUFA10 variants along with the NADH/NAD^+^ reporter SoNar. The NADH/NAD^+^ ratiometric image was constructed by taking the emission intensity of 405 nm (NADH binding) vs. 488 nm (NAD^+^ binding) for SoNar. Right, quantification of NADH/NAD^+^ ratio. (**F**) NDUFA10•C253D restores NADH oxidation upon reduced-state conditions. Flow cytometry analysis of the indicated NSCLC cells stably expressing SoNar and NDUFA10 variants (all PAM mutants) following depletion of endogenous NDUFA10. The NADH/NAD^+^ ratio was determined by comparing the emission intensity of 405 nm (NADH binding) vs. 488 nm (NAD^+^ binding) for SoNar. Heatmaps depict changes in mitochondrial NADH/NAD^+^ ratio relative to vehicle. Inset, representative NADH/NAD^+^ distribution for CALU6 cells under the indicated treatments. (**G**) NDUFA10•C253 exists as a sulfinic acid under basal conditions. Top, structure of the sulfinic acid probe NO-DTB. Bottom, lysates from CALU6 cell expressing the indicated proteins were treated with 500 µM NO-DTB. Sulfinic acid formation was determined following streptavidin-enrichment and immunoblot analysis. (**H**) Lowering ROS decreases NDUFA10 sulfinic acid modification. CALU6 cells were treated with 1 µM KI696, 5 µM DOXY for 48 hrs, or 10 mM NAC, 10 mM GSHee, and 250 µM Trolox for 1hr were incubated with 500 µM NO-DTB and sulfinic acid formation was determined as described in (**G**). (**I**) NRF2 activation alters NDUFA10 conformation in a C253-dependent manner. CALU6 cells depleted of endogenous NDUFA10 and expressing the indicated PAM mutants were treated with 1 µM KI696 or vehicle control for 48 hrs. Left, thermal stability analysis was conducted on corresponding cellular lysate by incubating at the indicated temperatures, followed by centrifugation and immunoblot analysis of NDUFA10 levels. Right, quantification of relative NDUFA10 levels. (**J**) Localized burst of H_2_O_2_ is sufficient to alter NDUFA10 conformation. CALU6 cells expressing FLAG-DAAO-NDUFA10 were treated with 10 mM L-Ala or D-Ala for 6 hrs, and thermal stability of NDUFA10 was determined by immunoblot (top) and quantified (bottom). Statistical significance was determined by Student’s *t* test (two-tailed, unpaired). ns, not significant; * *p* < 0.05; ** *p* < 0.01; *** *p* < 0.001. Scale bar: 25 µm.

These findings were particularly intriguing because higher-order cysteine oxidation states such as sulfinic acid modifications have been canonically associated with protein damage or inactivation (*40*). We therefore sought to directly test the sulfinic acid state of NDUFA10, using the NO-DTB probe, which selectively labels protein sulfinic acids (*24, 33*). We detected robust labeling of NDUFA10 that was lost in the C253S mutant (Fig. 4G). Across mouse tissues, NDUFA10•C253 oxidation varied, with brown adipose tissue showing the highest levels (fig. S6H). Decreasing cellular ROS levels further reduced NO-DTB labeling of NDUFA10•C253, indicating that NDUFA10 exists in a basally oxidized state which is reversible (Fig. 4H, fig. S6I).

Because our chemical proteomic analysis under different ROS states revealed both a decrease and increase in cysteine accessibility reflective of structural changes to the proteome, we next asked whether Cys253 oxidation alters NDUFA10 conformation. Thermal stability profiling (*78*) of NDUFA10 isolated from cells with NRF2 activation displayed reduced stability compared to controls, consistent with a conformational shift upon reduction (Fig. 4I). Conversely, H_2_O_2_ treatment stabilized NDUFA10 (fig. S6J). To determine whether local ROS production regulates NDUFA10 structure, we used the D-amino acid oxidase (DAAO) system to generate localized H_2_O_2_ bursts in response to D-Ala but not L-Ala (*79*) (fig. S6K, L). The NDUFA10-DAAO fusion exhibited D-Ala–dependent thermal stabilization, supporting a direct role for oxidation in regulating NDUFA10 conformation (Fig. 4J). Importantly, redox-induced stability shifts were absent in the C253D mutant under both reducing and oxidizing perturbations, implicating Cys253 as the dominant redox-sensitive site (Fig. 4I, J, fig. S6J). Collectively, these findings establish higher-order oxidation of a non-catalytic cysteine as a regulatory mechanism that links NDUFA10 redox conformational states to mitochondrial Complex I function.

### SRXN1 regulates NDUFA10 oxidation state

Having established the importance of oxidation of NDUFA10•C253 for Complex I function, we next sought to determine how this oxidation state is regulated during transitions between oxidized and reduced cellular environments. Cells rely on multiple oxidoreductase systems to buffer ROS, including superoxide dismutase (SOD), peroxiredoxins (PRDX), thioredoxin reductases (TXN), and sulfiredoxin (SRXN) (*25, 26, 80–82*), many of which are NRF2 targets (fig. S7A, B). We asked whether one or more of these systems controls the redox state of NDUFA10. Our CRISPR screens identified several oxidoreductases as modifiers of NRF2-induced cytotoxicity, with SRXN1 emerging as a top hit conferring resistance in CALU6 cells, a result we validated in additional NSCLC models (Fig. 5A, fig. S7C). Depletion of SRXN1 mitigated the NADH-reductive stress caused by NRF2 activation (fig. S7D–F), whereas SRXN1 overexpression increased NADH/NAD^+^ ratios and reduced Complex I activity (Fig. 5B, fig. S7G-J), suggesting that SRXN1 opposes oxidation-dependent NDUFA10 activity.

**Fig. 5:**
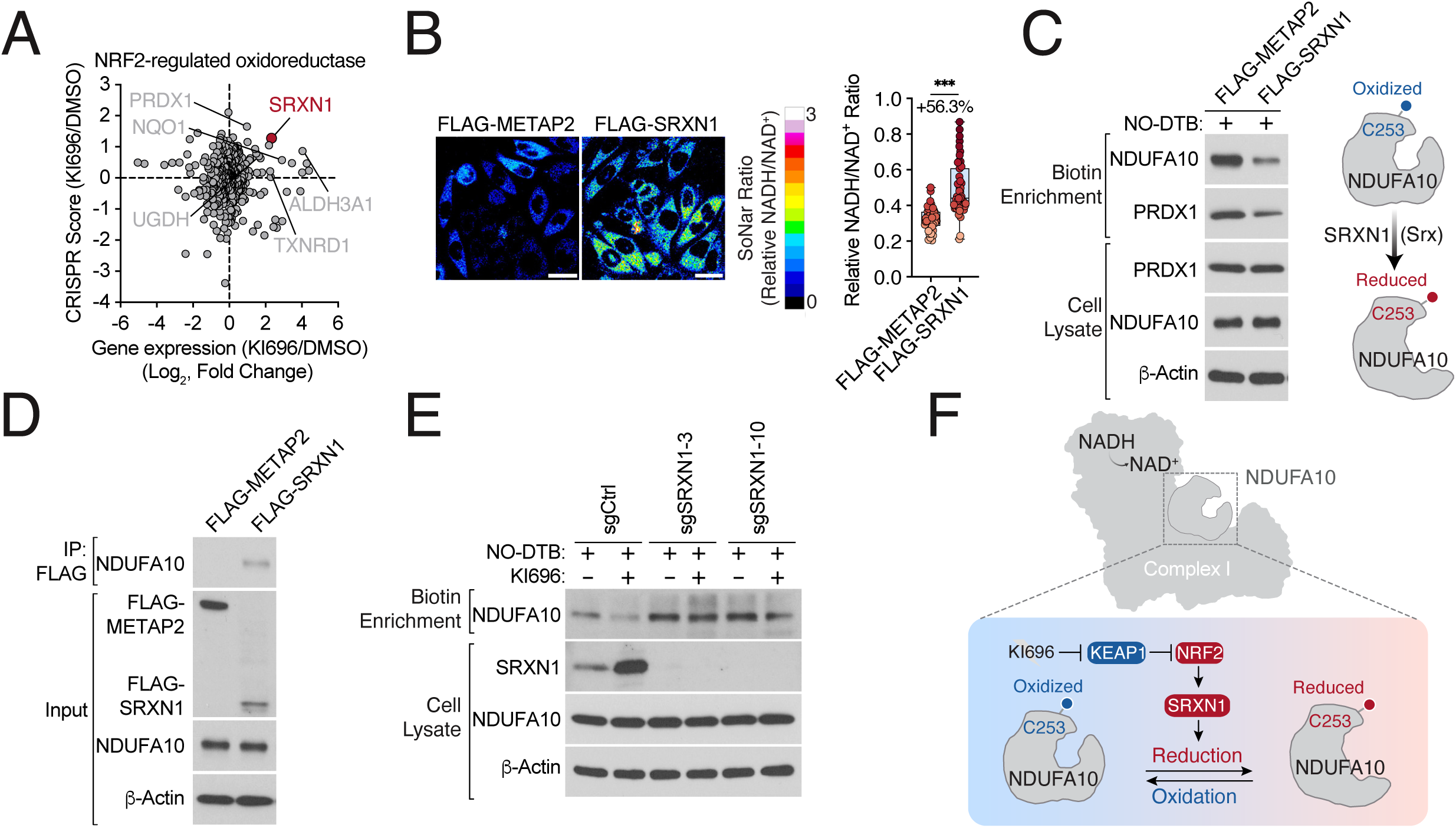
NRF2-SRXN1 axis controls NDUFA10 oxidation state. (**A**) Scatterplot comparing NRF2 target gene dependency and NRF2-regulated oxidoreductases identified in (*77*) in CALU6 cells. (**B**) SRXN1 over-expression results in a high NADH/NAD^+^ ratio. CALU6 cells stably expressing SoNar were transfected with METAP2 (control) or SRXN1, and the representative NADH/NAD^+^ ratio was assessed (left) and quantified (right). (**C**) SRXN1 regulates NDUFA10 sulfinylation. Left, over-expression of SRXN1 but not METAP2 (Control) in CALU6 cells decreases NO-DTB labeling of NDUFA10. Lysates were treated with 500 µM NO-DTB. Sulfinic acid formation was determined following streptavidin-enrichment and immunoblot analysis. PRDX1 serves as a positive control for SRXN1 activity. Right, schematic for SRXN1 mechanism of action. (**D**) SRXN1 interacts with NDUFA10. CALU6 cells expressing FLAG-METAP2 (Control) or FLAG-SRXN1 were lysed and the interaction between NDUFA10 and FLAG-tagged proteins was determined by immunoblot following immunoprecipitation from cell lysate with anti-FLAG M2 affinity beads. (**E**) SRXN1 regulates NDUFA10 oxidation. Depletion of SRXN1 increases NO-DTB labeling of NDUFA10 following NRF2 activation in CALU6 cells. NRF2 activation and NO-DTB labeling was assessed as described in (**C**). (**F**) Model, following NRF2 activation, SRXN1 reverses the sulfinylation of NDUFA10•C253. Statistical significance was determined by Student’s *t* test (two-tailed, unpaired). ns, not significant; * *p* < 0.05; ** *p* < 0.01; *** *p* < 0.001. Scale bar: 25 µm.

Although SRXN1 is classically characterized as an ATP-dependent reductase for hyperoxidized 2-Cys peroxiredoxins, chemical proteomic studies have suggested that sulfiredoxin-regulated activity may extend to additional protein substrates (*83*). We therefore asked whether SRXN1 regulates the sulfinic acid state of NDUFA10. Biochemically, SRXN1 overexpression decreased NO-DTB labeling of its canonical target PRDX1 along with NDUFA10, consistent with enhanced reversal of NDUFA10 sulfinylation in cells (Fig. 5C). SRXN1 contains a noncanonical mitochondrial targeting sequence and can translocate to mitochondria (*84, 85*), which we further confirmed in our models using immunofluorescence (fig. S8K) Immunoprecipitation assays further revealed a direct interaction between SRXN1 and NDUFA10 (Fig. 5D). Importantly, SRXN1 depletion restored sulfinic acid labeling of NDUFA10 following NRF2 activation (Fig. 5E). Because several reduced-state perturbations tested in this study cannot directly reduce sulfinic acids (*86*), we hypothesized that lowering intracellular ROS or increasing redox-buffering capacity would create a permissive environment for SRXN1 activity. Indeed, NAC treatment reduced NDUFA10 oxidation in cells overexpressing SRXN1 (fig. S7L, M). These results identify SRXN1 as a key regulator of the cellular NDUFA10 sulfinylation state and demonstrate that higher-order cysteine oxidation within Complex I is dynamically reversible through a major cellular redox control pathway (Fig. 5F).

### Oxidation-dependent nucleotide kinase activity of NDUFA10 controls mtDNA maintenance

Although NDUFA10 has been characterized primarily as a Complex I assembly factor, the marked conformational sensitivity of the protein to oxidation suggested that it might possess additional biochemical activities regulated by redox state that support complex I function. We therefore sought to determine whether oxidation-dependent structural changes in NDUFA10 directly alter its biochemical activity.

Sequence and structural analysis revealed a conserved nucleotide-binding pocket within NDUFA10, peripheral to Cys253 (Fig. 6A). Although NDUFA10 contains a deoxy nucleotide kinase (dNK) domain and shares strong catalytic sequence and structural similarity with canonical dNK enzymes, particularly dGK, it lacks detectable dGK activity (Fig. 6A, fig. S8A, B). Examination of solved structures revealed electron density consistent with bound nucleotides, including dGTP, ATP, or ADP (Fig. 6B, fig. S8C) (*87–91*), with the former having been recently demonstrated to interact with NDUFA10 (*88*). We further identified residues predicted to coordinate magnesium and stabilize phosphate groups, suggesting a potential nucleotide kinase reaction: ATP + dGDP ⇌ ADP + dGTP (Fig. 6A). To test this, we purified WT NDUFA10, the reduced mutant (C253S), and a magnesium binding mutant (E88A), and measured nucleotide conversion by HPLC (fig. S8D). WT NDUFA10 catalyzed phosphate transfer from ATP to dGDP, generating dGTP (Fig. 6C). This activity was absent in the magnesium-binding mutant and markedly attenuated in the reduced C253S mutant (Fig. 6C). Addition of EDTA blocked kinase activity indicating the need for divalent metal ions, and we found that the reaction was reversible, with dGTP serving as phosphate donor in the reverse reaction (fig. S8E).

**Fig. 6:**
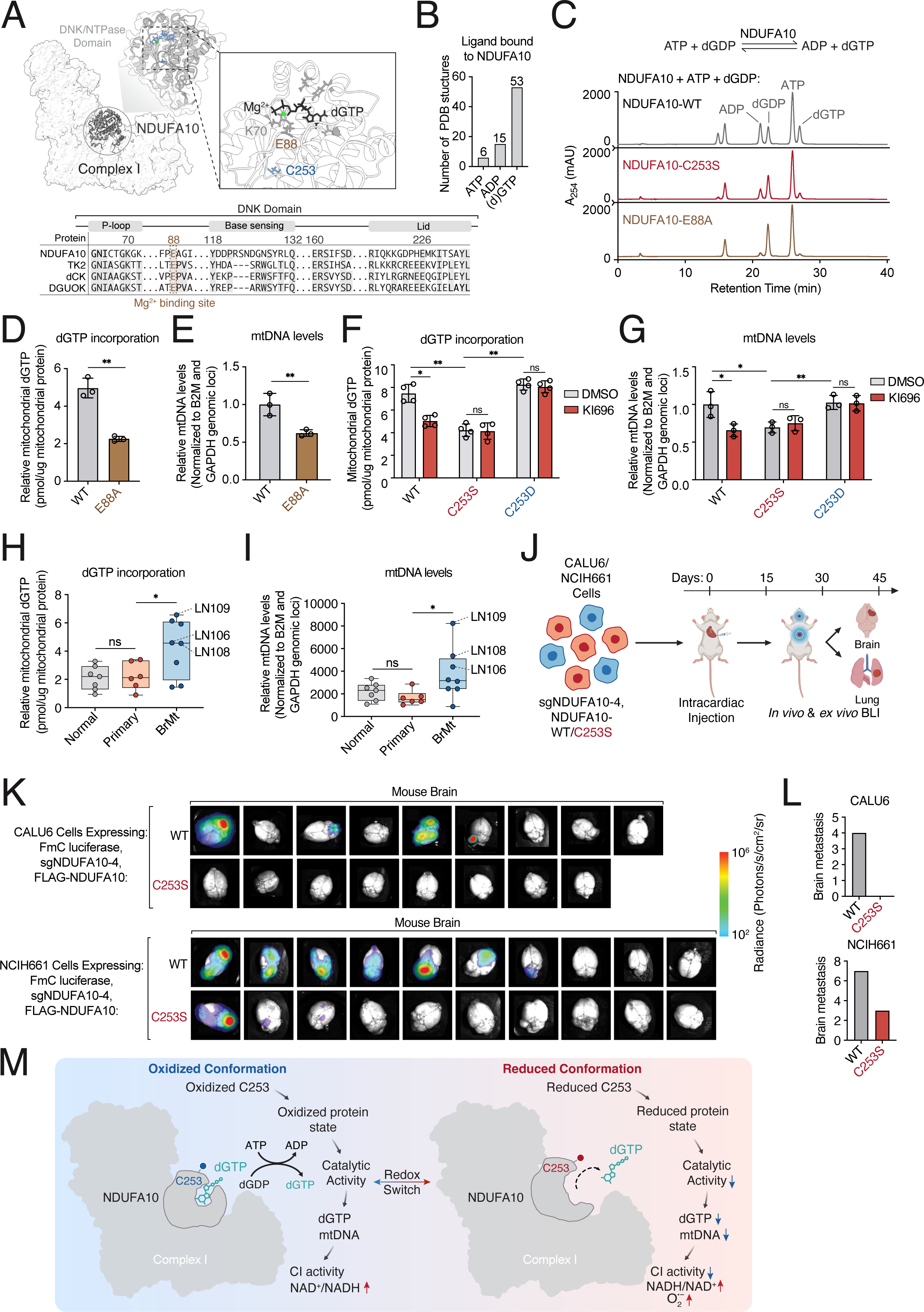
NDUFA10 oxidation gates nucleotide kinase activity, mtDNA maintenance, and brain metastases. (**A**) NDUFA10 has a putative deoxy nucleotide kinase (DNK) domain. Left, location of NDUFA10 in Complex I (PDB:9TI4). Middle, enhanced view of putative NDUFA10 DNK-like domain (dark grey) highlighting the relative position of C253 proximal to the nucleotide binding pocket. Right, Ligand-bound structure of NDUFA10 highlighting electron densities corresponding to dGTP and Mg^2+^ and corresponding residues important for catalysis. Bottom, sequence alignment of NDUFA10 with DNK domains highlighting conserved features required for catalysis (see also Methods). (**B**) Quantification of putative nucleotides bound to NDUFA10 in resolved PDB structures (see also fig. S9C). (**C**) NDUFA10 is a redox-gated dGTP kinase. Representative HPLC chromatograms (n=3) showing dGTP synthesis following incubation with 1.5 µM of recombinant NDUFA10-WT, C253S (reduced mimetic) or E88A (kinase dead) variants (see also fig. S9D, E, Methods). (**D**, **E**) NDUFA10 regulates mitochondrial dGTP incorporation and mtDNA levels. Relative mitochondrial dGTP incorporation into mtDNA (D) and mtDNA (E) were measured in CALU6 cells depleted of endogenous NDUFA10 and expressing PAM-mutants of NDUFA10 WT or its kinase-dead variant (E88A). dGTP incorporated into mtDNA was measured by a modified qPCR assay (see also fig. S10C, Methods). mtDNA levels were quantified qPCR of MT-ND1 and MT-CO1 and normalized to the nuclear genes B2M and GAPDH. (**F, G**) Cellular NDUFA10 kinase activity is oxidation-dependent. Mitochondrial dGTP (F) and mtDNA levels (**G**) were quantified as described in (**D, E**) from CALU6 cells expressing the indicated NDUFA10 variants in NDUFA10-depleted background following treatment with 1 µM KI696 or vehicle control for 48 hrs. (**H, I**) Brain metastases contain elevated mitochondrial dGTP and mtDNA relative to primary tumors and normal lung. Mitochondrial dGTP (**H**) and mtDNA (**I**) were quantified from tumors and normal lung tissue as described in (**D, E**). **(J**) Schematic, testing the requirement of NDUFA10 oxidation in metastatic fitness in the brain. Cells co-expressing sgNDUFA10 together with FLAG-tagged WT or C253S NDUFA10 PAM mutants and luciferase were injected intracardiacally into mice, followed by *in vivo* and *ex vivo* bioluminescence imaging to quantify tumor burden. (**K, L**) NDUFA10 oxidation supports brain metastases. Left, representative *ex vivo* bioluminescence images (BLI) of mouse brains isolated from intracardiac injection of CALU6 and NCIH661 cells depleted of endogenous NDUFA10 and expressing the indicated mutants (**K**). Right, quantification of brain metastatic burden (CALU6-NDUFA10-WT: n=9; NDUFA10•C253S: n=8; NCIH661-NDUFA10-WT: n=10; NDUFA10•C253S: n=10) (**L**). (**M**) Model, oxidation of NDUFA10•C253 promotes dGTP kinase activity, leading to mtDNA maintenance and Complex I function, and support lung cancer metastasis to the brain. Statistical significance was determined by Student’s *t* test (two-tailed, unpaired). ns, not significant; * *p* < 0.05; ** *p* < 0.01; *** *p* < 0.001.

We next asked whether oxidation directly regulates NDUFA10 kinase activity. Addition of H_2_O_2_ enhanced kinase activity in a dose-dependent manner, an effect lost in the C253S mutant (fig. S8F, G), indicating that oxidation promotes a conformation permissive for catalysis. Consistent with this, a biotinylated dGTP analog bound more strongly to the oxidized-mimetic C253D mutant than to the reduced mutant (fig. S9A). Furthermore, addition of dGTP stabilized NDUFA10 and its interacting partners in lysate (fig. S9B), consistent with product-induced conformational stabilization. Importantly, expression of the kinase-dead E88A mutant also increased the NADH/NAD⁺ ratio (fig. S9C), connecting NDUFA10 nucleotide kinase activity to Complex I function.

Because mitochondrial dGTP availability is required for mtDNA replication and maintenance, we next examined whether NDUFA10 redox state regulates mitochondrial nucleotide pools in cells. Using a qPCR-based assay to measure dGTP incorporation into mtDNA (fig. S9D), we found that expression of either NDUFA10•C253S or the kinase-dead E88A mutant in NDUFA10-depleted cells, markedly reduced dGTP levels (Fig. 6D, F, fig. S9E, F). NRF2 activation similarly reduced dGTP incorporation, which was completely rescued in cells expressing the oxidation-mimetic NDUFA10•C253D mutant (Fig. 6F, fig. S9F). Consistent with a decrease in dGTP levels, we found a concomitant reduction in mtDNA in cells expressing the NDUFA10 kinase dead or reduced mutant (Fig. 6E, G, fig. S9G), providing an explanation for the loss of Complex I function. Collectively, these results identify a previously unrecognized nucleotide kinase activity for NDUFA10 that is gated by oxidation state and couples the mitochondrial redox environment to nucleotide metabolism, mtDNA maintenance and complex I function.

### NDUFA10 oxidation supports brain metastasis

Our patient analyses demonstrated that brain metastases preferentially occupy oxidized protein states, and our mechanistic studies identified oxidation of NDUFA10 as a key regulator of Complex I function and mitochondrial nucleotide metabolism. We therefore asked whether this oxidation-dependent pathway contributes directly to metastatic fitness *in vivo*.

To first determine whether the pathway is active in human tumors, we quantified dGTP incorporation into mtDNA across patient samples. Brain metastases displayed markedly higher dGTP incorporation and mtDNA levels than primary tumors when normalized to adjacent normal tissue (Fig. 6H, I, fig. S9H), consistent with prior analyses of paired primary lung tumors and brain metastases showing increased mtDNA content and mitochondrial features in metastatic lesions (*92*). Prior studies have implicated Complex I in metastatic fitness across multiple cancers (*93–95*). Given that brain metastases exhibited oxidized protein states, the mitochondria in these tumors was enriched for oxidized protein states, and that NDUFA10 oxidation supports Complex I activity and mitochondrial nucleotide pools, we hypothesized that oxidation of NDUFA10 may be specifically required for metastatic colonization. Expression of the reduced-state mutant NDUFA10•C253S impaired migration and invasion *in vitro* in four NSCLC models (fig. S10A), suggesting a role in metastatic competence. To test this directly *in vivo*, we generated CALU6 and NCIH661 models expressing WT or reduced-state NDUFA10 in the context of endogenous NDUFA10 depletion and labeled cells with luciferase for bioluminescent imaging (Fig. 6J). Following intracardiac injection, both WT and reduced-mutant cells formed lung lesions with similar efficiency (fig. S10B-G). In contrast, WT cells efficiently colonized the brain, whereas cells expressing the reduced-state mutant showed a striking reduction in brain metastases (Fig. 6K, L). To isolate colonization effects from dissemination, we performed intracranial injections with the NCIH661 cell line. Tumors expressing the NDUFA10 reduced-state mutant exhibited significantly impaired growth relative to WT controls (fig. S10H). Together, these results reveal that high-order cysteine oxidation of NDUFA10 supports metastatic fitness within the brain, and provide a mechanistic link between oxidized protein states, mtDNA levels, mitochondrial Complex I function, and metastatic progression (Fig. 6M).

## Discussion

A longstanding challenge in cancer biology has been to understand how the tumor environment shapes cellular behavior beyond what can be inferred from genetic or transcriptional states (*16, 96*). Redox biology exemplifies this gap: although regulation of oxidative stress has been ubiquitously linked to tumor initiation, growth, and metastasis (*2–4*), the field has lacked a systematic framework that connects variation in the ROS redox state to defined molecular mechanisms and cancer phenotypes. In this study, we demonstrate that ROS-associated protein states can be captured in human tumors at the level of cysteine accessibility, providing a structural and mechanistic complement to genetic and transcriptomic descriptions of tumor cell states. This work further supports the premise that protein oxidation can function as a regulatory input that shapes tumor fitness, rather than merely a source of molecular damage.

Cysteine accessibility is directly influenced by the intracellular environment. Accordingly, chemical proteomics allows us to understand the corresponding ensemble of protein states under a given environmental condition. By defining signatures of tumor redox environments through changes in cysteine accessibility, this work reframes redox biology through a structural lens. This is particularly relevant in metastasis, where cancer cells encounter extreme microenvironmental constraints and where redox sensitivity has been repeatedly observed but poorly understood at the level of proteome (*11, 13, 16*). Thus, chemical proteomic signatures provide a residue-level connection linking changes in the cellular environment to protein function and pathways that support cancer progression. In this context, we were surprised to find that many active tumors displayed oxidative protein states, given the prevailing view that highly oxidized environments are detrimental to tumor fitness (*7, 11, 13*). This discrepancy suggests that transcriptomic and protein-state measurements capture distinct layers of biological regulation that do not necessarily always align. For example, activation of an antioxidant transcriptional program may represent a compensatory response to oxidative stress within the tumor microenvironment, while proteins remain oxidized at the post-translational level.

Combining our analysis of ROS-associated protein states with functional genomics helps explain how redox pressures translate into selective advantages during metastatic colonization. Importantly, we uncovered how an oxidizing protein state can support mitochondrial Complex I function rather than simply acting as non-specific stress. Because cysteine accessibility integrates multiple biochemical inputs, including cysteine oxidation and conformational changes (*35, 36, 53*), future studies that more precisely disentangle these contributions across different cellular environments will be essential for defining the structural landscape of cells, tumors, and organisms using chemical proteomic technologies.

High levels of oxidative stress are damaging to the electron transport chain (*73, 97–99*). Our findings extend these observations by demonstrating that Complex I function depends on a balance of cysteine redox states, including site-specific oxidation. Like other metabolites, ROS must be tightly regulated to achieve cellular homeostasis (*3, 100, 101*). Accordingly, cysteine oxidation and reduction must be tailored to specific cellular contexts. Our results begin to suggest that mitochondrial H_2_O_2_ is more than a byproduct of Complex I activity, but rather a necessary regulatory component (*19, 102, 103*) sensed by cysteines embedded in this complex. The finding that sulfinic acid modification of NDUFA10 promotes a functional NDUFA10 state is conceptually consistent with other studies which demonstrate that negatively charged protein modifications, including phosphorylation, function as allosteric modulators (*104, 105*). The identification that oxidation at a specific, non-catalytic cysteine gates NDUFA10 nucleotide kinase activity to support mitochondrial genome maintenance, provides a clear example of this logic. Accordingly, a decrease in mtDNA levels underlies the decrease in Complex I function (*48*) observed in cells expressing constitutively reduced NDUFA10. This work supports the premise that protein oxidation can function as a regulatory input at the level of protein conformations rather than non-specific damage, and help explains why perturbations that suppress ROS can create unanticipated vulnerabilities. Moreover, our findings additionally provide a mechanistic link between ROS-mediated damage of mtDNA and pathways that replenish nucleotide pools. Thus, the presence of a ROS-sensitive cysteine embedded within Complex I may serve to preserve the integrity of mitochondrial genome following oxidative nucleotide damage generated during normal mitochondrial activity.

In conclusion, we provide a biochemical landscape of oxidized and reduced protein states in human lung cancers and demonstrate that a subset of lung tumors, particularly brain metastases, are characterized by oxidized protein states. We identify a ROS-sensitive allosteric site embedded within mitochondrial Complex I whose high-order cysteine oxidation supports Complex I function through mtDNA regulation, linking ROS states with mitochondrial genome maintenance through nucleotide production. More broadly, these findings establish ROS-associated protein conformations as a functional dimension of tumor progression and suggest that therapeutic strategies that selectively control cysteine oxidation may provide new opportunities to target metastatic lesions.

## Acknowledgements

We thank all members of the Bar-Peled and Brastianos lab for helpful suggestions. This work was supported by the Damon Runyon Cancer Research Foundation (62-20 to L-B.P., B.L.L.), MGH ECOR Fund for Medical Discovery (to M.Ge), the American Association for Cancer Research (19-20-45-BARP to L. B-P., 22-30-73-BRAS to P.K.B ), the American Cancer Society, the Melanoma Research Alliance (to L.B,-P, P.K.B), the Ludwig Cancer Center of Harvard Medical School, Lungevity, ALK Positive, V-Foundation, the Mark Foundation, Mary Kay Foundation, Paula and Rodger Riney Foundation, the PEW-Stewart Trusts, Mark Foundation, Lisa and Mark Schwartz, Carol and Ben Monderer, Eileen and Jim Rullo, JST SPRING, Japan (JPMJSP2123 to Y.S.), the Toshizo Watanabe Foundation (Y.S.), and support from the Yamagata prefectural government and Tsuruoka City, Japan (Y.S.), the NIH/NCI (K99CA296772 to M.Ge, 1R21CA226082-01, R37CA260062, R01CA285415 to L.B-P., 5R01CA227156, 1R01CA294793 to P.K.B., 2T32HG010464-06 to C.T.), the NIH/NHLBI (R01HL177625 to Q.L.), Krantz Breakthrough Award (to M.L.S. and P.K.B.), Philanthropic Award from Alexandra Drane (to P.K.B.), Breast Cancer Research Foundation (to P.K.B), National Brain Tumor Society (to P.K.B.), Terry and Jean de Gunzberg MGH Research Scholar Award (to P.K.B.), and the Hellenic Women’s Club (to P.K.B.). E.T.C is an HHMI Investigator.

## Author Contributions

L.B.-P., P.K.B. and M.Ge conceived and designed the study. M.Ge performed most of experiments with assistance from M.Gohar, Y.S., S.J., Y.H., B.W., J.W., M.Y.A., S.H., C.Cakici, Z.D., N.V., Z.Q., J.Z., I.Z., C.Che, R.S., N.K., J.J., Y.X., S.I., C.L., Q.L., F.-Y.W., M.L.S., B.B.L., A.N.H., E.T.C. and A.J.I.. C.T. and N.N. performed mouse experiments. M.Gohar performed bioinformatics analysis with assistance of H.F., P.H., M.S.L. and D.G.. E.S., C.W.M., M.M.-L. and L.S. assisted with patient clinical information and cohort annotation. M.Ge, M.Gohar, C.T., Y.S, P.K.B. and L.B.-P. wrote the manuscript with assistance from all the coauthors. L.B.-P. supervised the studies.

---

**Fig. S1:**
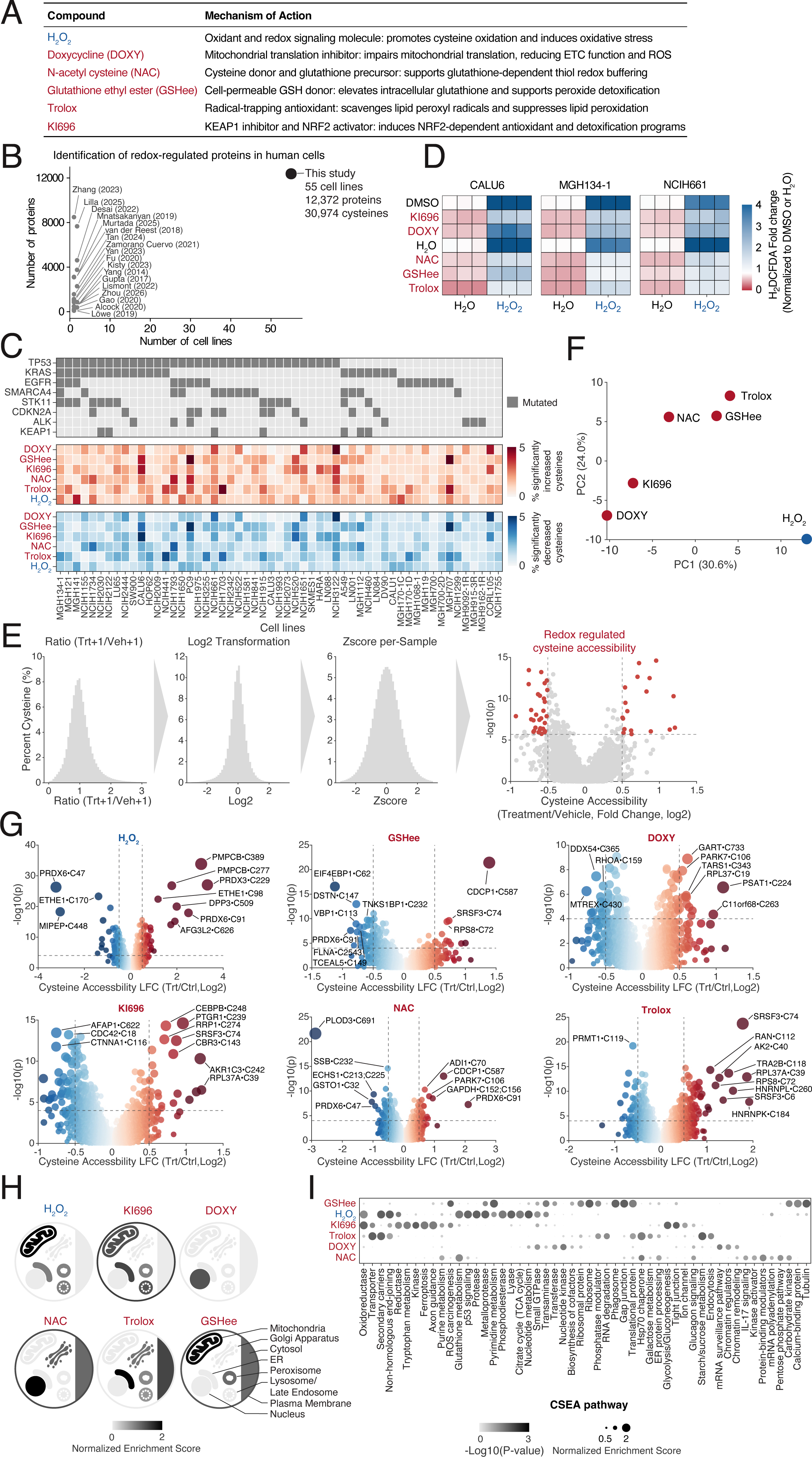
Characterization of cysteines impacted by ROS perturbations in NSCLCs. (**A**) Table summarizing ROS perturbations used in this study and proposed mechanisms of action. (**B**) Comparison of recent chemical proteomic studies of redox regulation in human cell lines. (**C**) Heatmap summarizing cysteine accessibility changes across NSCLC models following ROS perturbations. (**D**) Cellular antioxidant programs and redox-state perturbations alter intracellular ROS levels. The indicated NSCLC cell lines were treated with 1 µM KI696 or 5 µM DOXY for 48 hrs, or 10 mM NAC, 10 mM GSHee, 250 µM Trolox for 1 hr, and ROS levels were determined by flow cytometry using H_2_DCFDA probe following 100 µM H_2_O_2_ treatment for 30 min. (**E**) Schematic for normalization of cysteine accessibility (see also Methods). (**F**) PCA analysis of cysteine accessibility changes following different redox perturbations. (**G**) Defining redox-associated cysteine accessibility changes. Volcano plot depicting statistically significant changes in cysteine accessibility across treatments in this study. Median accessibility changes were reported across the NSCLC models analyzed. (**H**) Cellular compartments impacted by redox. Cellular map showing normalized enrichment score (NES) following each perturbation (also see Methods). (**I**) Cellular pathways impacted by redox. CSEA enrichment scores for pathways with cysteine accessibility differences following each redox treatment.

**Fig. S2:**
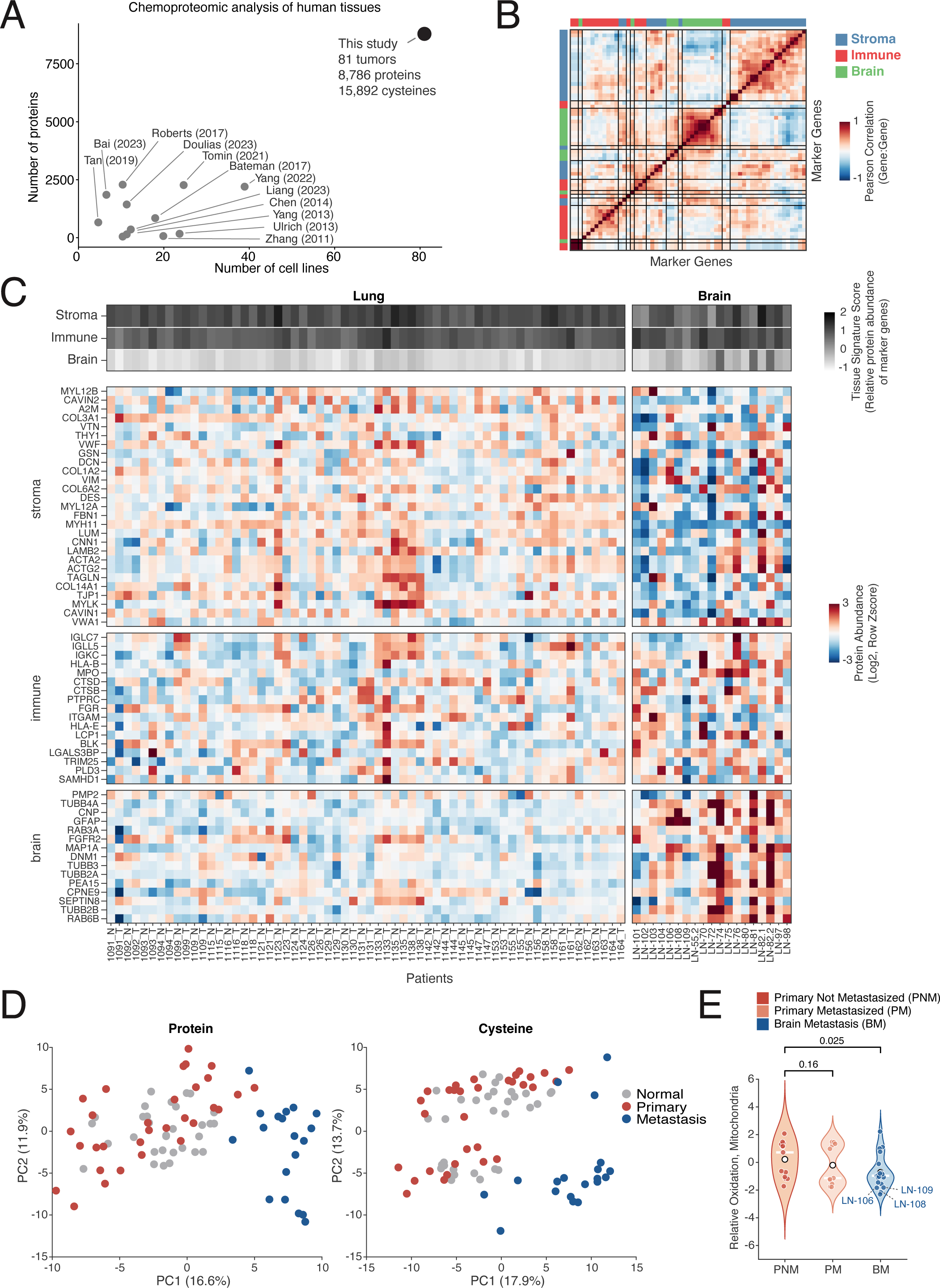
Mapping cysteine accessibility changes in human lung tumors. (**A**) Comparison of recent chemical proteomic studies of redox regulation in human tissues. (**B**) Correlation heatmap of tissue signature scores among specimens profiled. (**C**) Tissues signature scores. Top, composite tissue signatures for each specimen analyzed. Bottom, heatmap depicting relative expression of marker proteins for stromal, epithelial and brain tissues in each specimen (also see methods). (**D**) Brain metastases are distinct from normal and malignant lung tissue. PCA clustering of specimens by protein abundance (left) and cysteine accessibility (right). (**E**) Metastatic primaries and brain metastases have oxidative mitochondria relative to non-metastatic primaries (one-tailed Wilcoxon rank-sum test).

**fig. S3:**
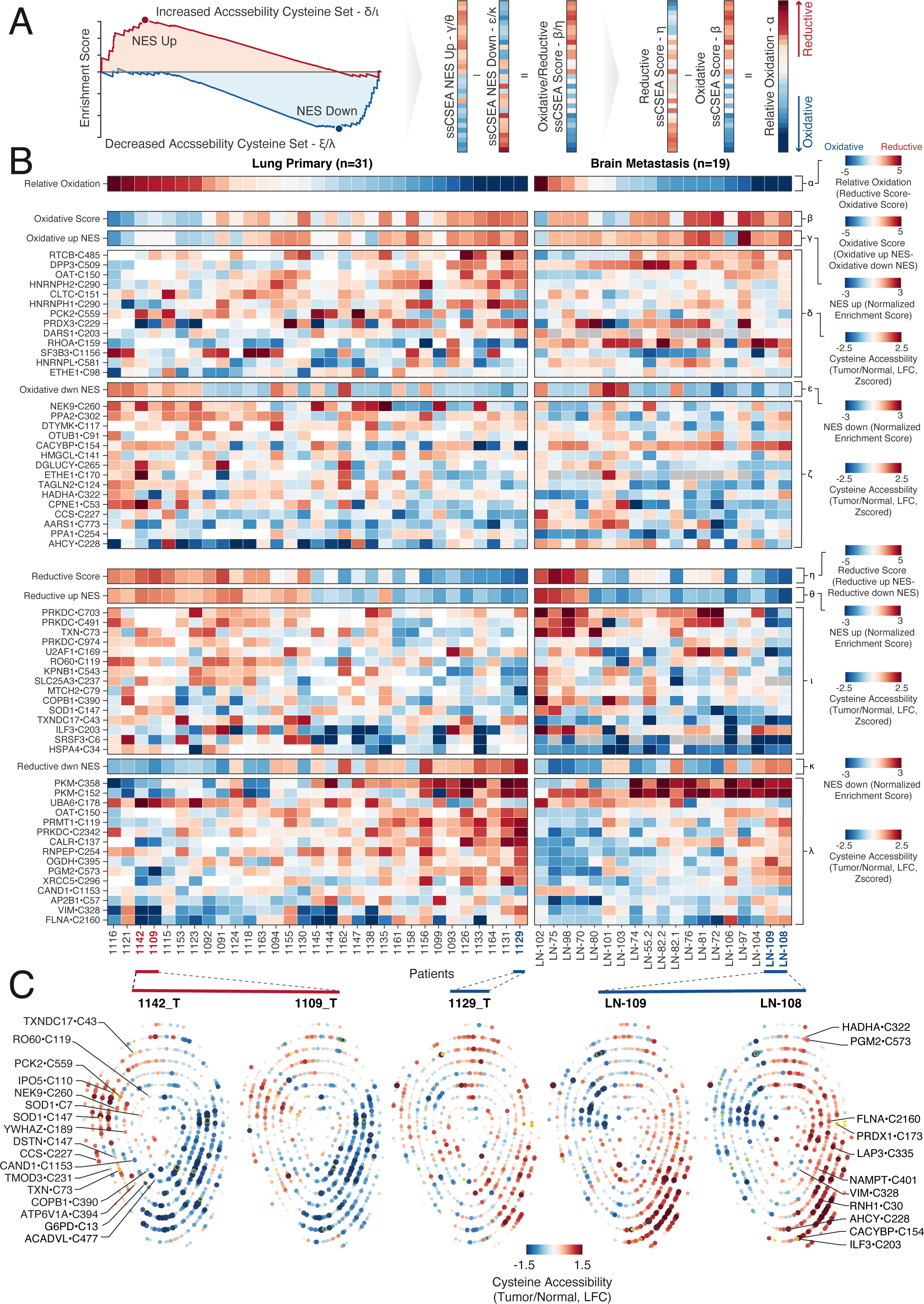
Differential redox-associate protein states in lung primary tumors and brain metastases. (**A**) Schematic of relative oxidation score derivation. For each signature, increased- and decreased-accessibility cysteine sets are independently scored by single-sample cysteine enrichment analysis (ssCSEA). Reductive and oxidative composite scores are then calculated from their corresponding signatures, and relative oxidation is determined as the difference between reductive and oxidative scores (see also Methods). **(B)** Distribution of reductive and oxidative protein states in human lung tumors and brain metastases. Heatmaps depict select cysteine accessibility changes comprising oxidative and reductive redox-state signatures, alongside corresponding ssCSEA enrichment and composite scores for each specimen (see also Methods). (**C**) Fingerprint plots taken from specimens with higher levels of reduced protein states (red) or oxidized (protein states).

**Fig. S4:**
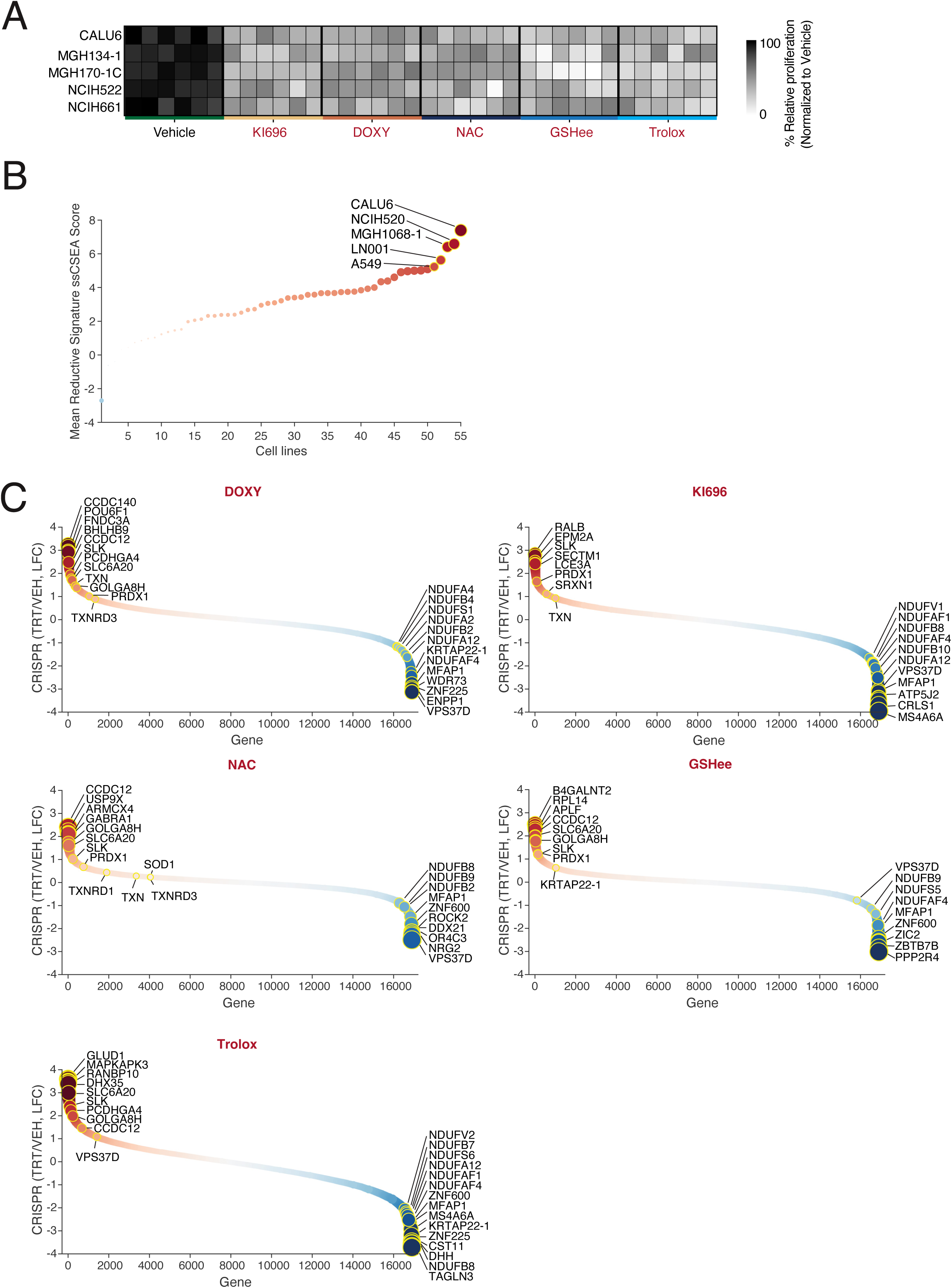
Functional genomic identification of genes that mediate sensitivity or resistance to lowering ROS levels. (**A**) Decreasing ROS levels slows NSCLC proliferation. The indicated cell lines were treated with 1 µM KI696, 5 µM DOXY, 10 mM NAC, 10 mM GSHee, 250 µM Trolox, and cellular proliferation was determined at 120 hrs by measuring relative DNA levels using Hoechst 33342. (**B**) Reductive signature in NSCLC cell lines under reduced-state conditions. (**C**) Identification of genes that mediate sensitivity or resistance to perturbations that lower ROS levels. Scatter plots depicting gene-level CRISPR scores in CALU6 cells following each treatment.

**fig. S5:**
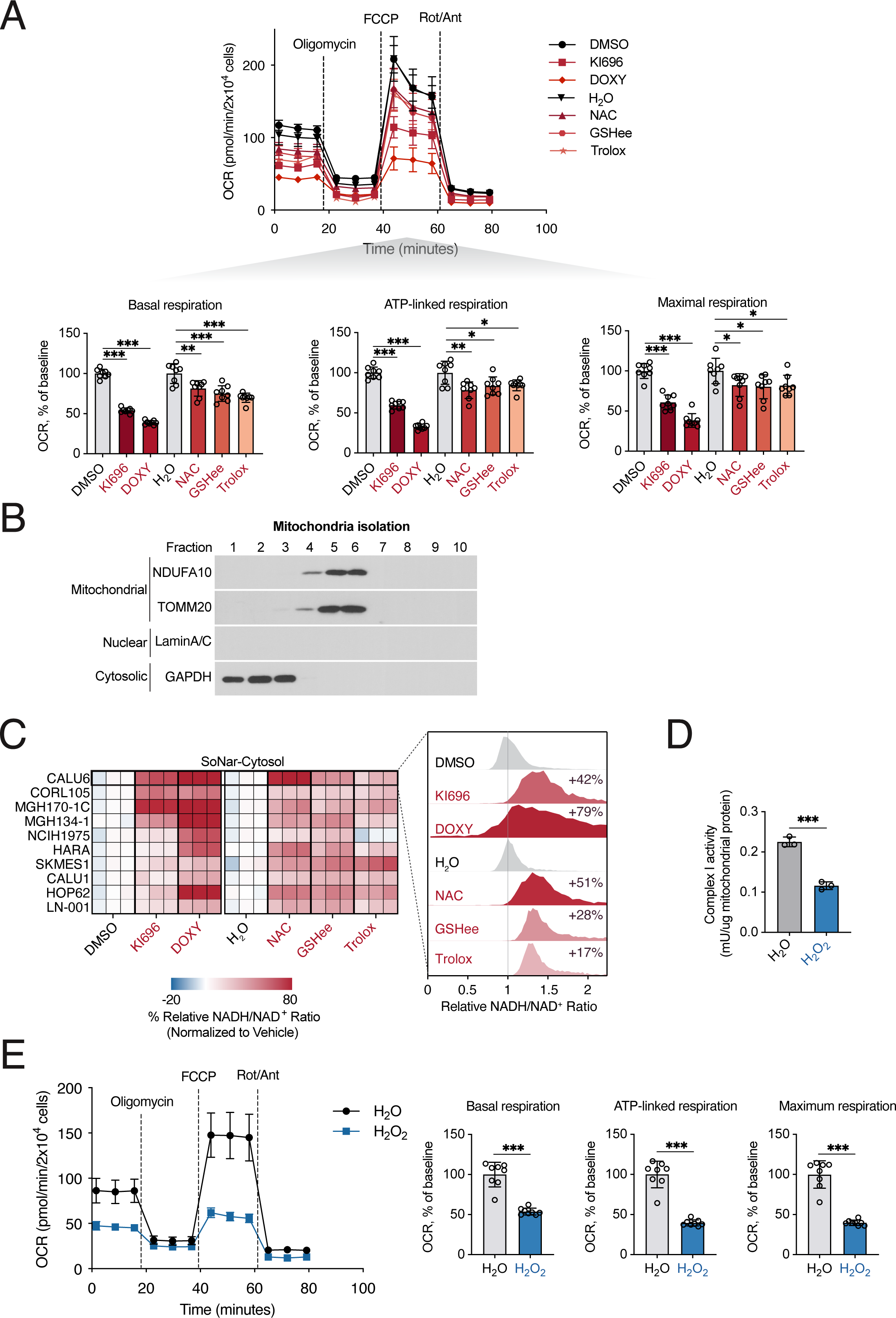
Redox perturbations impair mitochondrial function. (**A**) Reduced cell state impairs mitochondrial respiration. MGH134-1 cells were treated with 1 µM KI696 or 5 µM DOXY for 48hrs, or 10 mM NAC, 10 mM GSHee, and 250 µM Trolox for 1 hr, and oxygen consumption rate (OCR) was measured by Seahorse analysis. Bottom, quantification of basal respiration, maximal respiration, and ATP-linked respiration from Seahorse experiments. (**B**) Mitochondrial fractionation. Immunoblot analysis of the indicated fractions showing enrichment of NDUFA10 and TOMM20 in mitochondrial fractions, with LaminA/C and GAPDH marking nuclear and cytosolic fractions, respectively. (**C**) Reduced cell states increase the cytosolic NADH/NAD^+^ ratio. Cells were treated with the indicated compounds and cytosolic NADH/NAD^+^ was measured using SoNar localized in the cytosol. Left, heatmap depicts percent change in the cytosolic NADH/NAD^+^ ratio across NSCLC cell lines relative to vehicle. Right, representative distribution plots from CALU6 cells. The NADH/NAD^+^ ratio was determined by comparing the emission intensity of 405 nm (NADH binding) vs. 488 nm (NAD^+^ binding) for SoNar. (**D**) H_2_O_2_ decreases mitochondrial Complex I activity. CALU6 cells were treated with H_2_O or 100 µM H_2_O_2_ for 1 hr and Complex I activity was measured in isolated mitochondria following the addition of NADH and ubiquinone_1_ and normalized to mitochondrial protein abundance. (**E**) Acute H_2_O_2_ treatment suppresses mitochondrial respiration. Left, representative Seahorse traces in CALU6 cells treated with H_2_O or 100 µM H_2_O_2_ for 1 hr. Right, quantification of basal respiration, maximal respiration, and ATP-linked respiration. Statistical significance was determined by Student’s *t* test (two-tailed, unpaired). * *p* < 0.05; ** *p* < 0.01; *** *p* < 0.001.

**Fig. S6:**
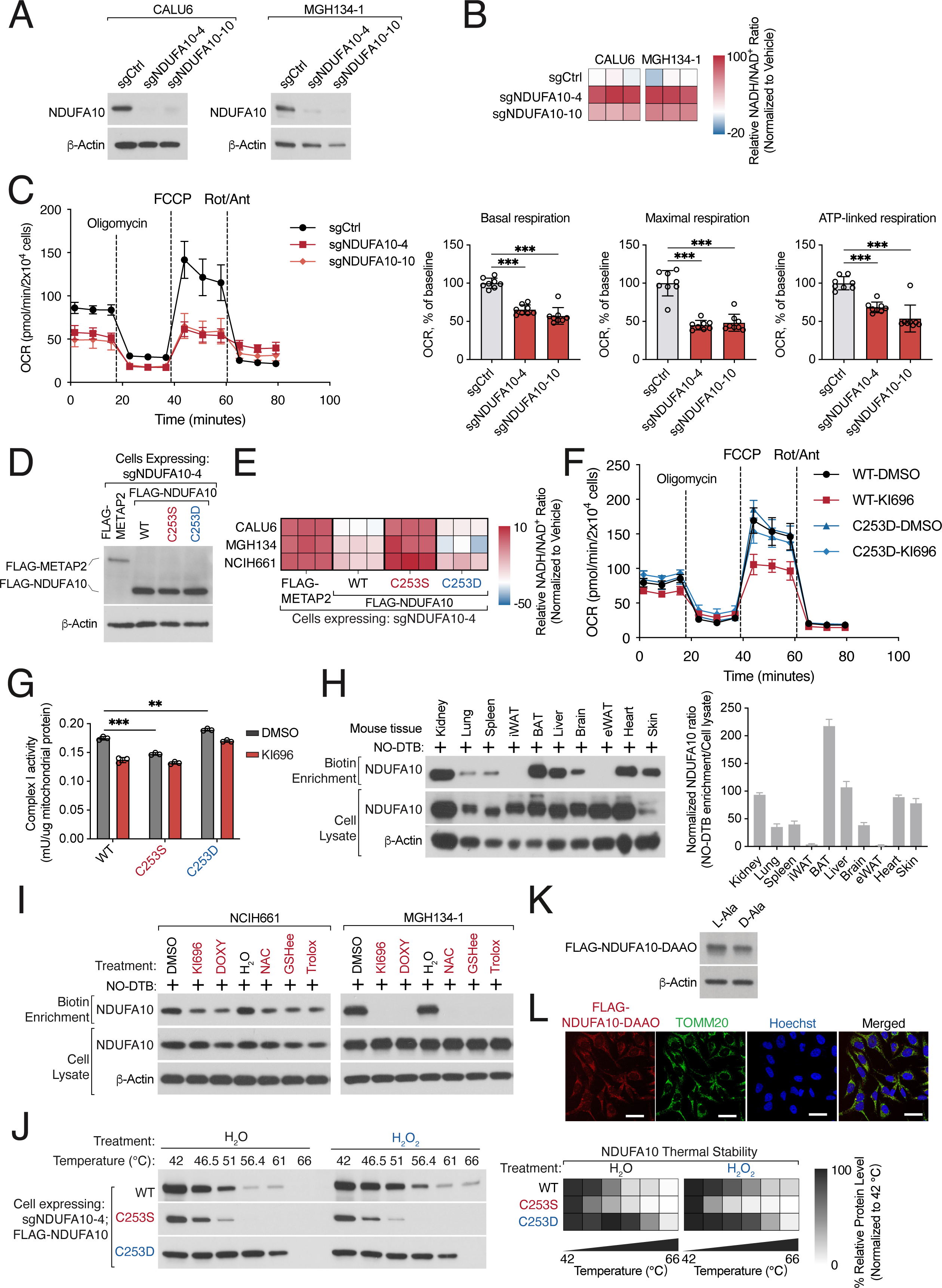
Cysteine oxidation of NDUFA10 regulates its conformation and Complex I activity. (**A**) Immunoblot validation of NDUFA10 depletion in CALU6 and MGH134-1 cells expressing the indicated sgRNAs. (**B**) Depletion of NDUFA10 increases the NADH/NAD^+^ ratio. Flow cytometry analysis of NSCLC cells stably co-expressing SoNar and the indicated sgRNAs targeting NDUFA10. The NADH/NAD^+^ ratio was determined by comparing the emission intensity of 405 nm (NADH binding) vs. 488 nm (NAD^+^ binding) for SoNar. (**C**) Loss of NDUFA10 impairs mitochondrial respiration. Left, representative Seahorse traces of oxygen consumption rate (OCR) in CALU6 cells expressing control sgRNA or sgRNAs targeting NDUFA10. Right, quantification of basal respiration, maximal respiration, and ATP-linked respiration. (**D**) Immunoblot analysis of the indicated proteins (all PAM mutants, except METAP2) in the background of NDUFA10 depleted CALU6 cells. (**E**) The NDUFA10 reduced mimetic (NDUFA10•C253S) has a defect in maintenance of NADH/NAD^+^ ratio. Flow cytometry analysis of the indicated NSCLC cells stably expressing SoNar and the indicated proteins (all PAM mutants) following depletion of endogenous NDUFA10. The NADH/NAD^+^ ratio was determined by comparing the emission intensity of 405 nm (NADH binding) vs. 488 nm (NAD^+^ binding) for SoNar. (**F**) The oxidized-state mimetic NDUFA10•C253D partially preserves mitochondrial respiration under KI696 treatment. Oxygen consumption in CALU6 cells co-expressing sgNDUFA10_4 and PAM-mutants of NDUFA10 WT or C253D using Seahorse bioanalyzer. (**G**) NDUFA10 oxidation partially restores Complex I activity following NRF2 activation. CALU6 cells co-expressing sgNDUFA10_4 and PAM-mutants of NDUFA10 WT, C253S or C253D were subjected to mitochondrial isolation and Complex I activity was determined following the addition of NADH and ubiquinone_1_. (**H**) NDUFA10 sulfinic acid oxidation is detected across mouse tissues. Left, NO-DTB enrichment of sulfinylated NDUFA10 from the indicated mouse tissues followed by immunoblotting. Right, quantification of enriched NDUFA10 normalized to input NDUFA10 signal. (**I**) Lowering ROS decreases NDUFA10 sulfinic acid oxidation in NSCLC cell lines. Lysates from NCIH661 and MGH134-1 cells previously treated with 1 µM KI696 or 5 µM DOXY for 48 hrs, or 10 mM NAC, 10 mM GSHee, 250 µM Trolox for 1 hr were incubated with 500 µM NO-DTB and sulfinic acid formation was determined following streptavidin-enrichment and immunoblot analysis. (**J**) H_2_O_2_ increases the thermal stability of NDUFA10. CALU6 cells co-expressing sgNDUFA10_4 and indicated PAM mutants of NDUFA10 WT, C253S, or C253D mutant, were lysed and treated with 100 µM H_2_O_2_ for 1 hr, incubated at the indicated temperatures, followed by centrifugation. Relative NDUFA10 levels were analyzed by immunoblot (left) and quantified (right). (**K, L**) Immunoblot analysis of CALU6 cells expressing FLAG-DAAO-NDUFA10 treated with 10 mM L-Ala or D-Ala for 6 hrs (**K**). Representative immunofluorescence images demonstrating co-localization of FLAG-DAAO-NDUFA10 with mitochondrial markers in CALU6 cells (**L**). Statistical significance was determined by Student’s *t* test (two-tailed, unpaired). ns, not significant; ** *p* < 0.01; *** *p* < 0.001.

**Fig. S7:**
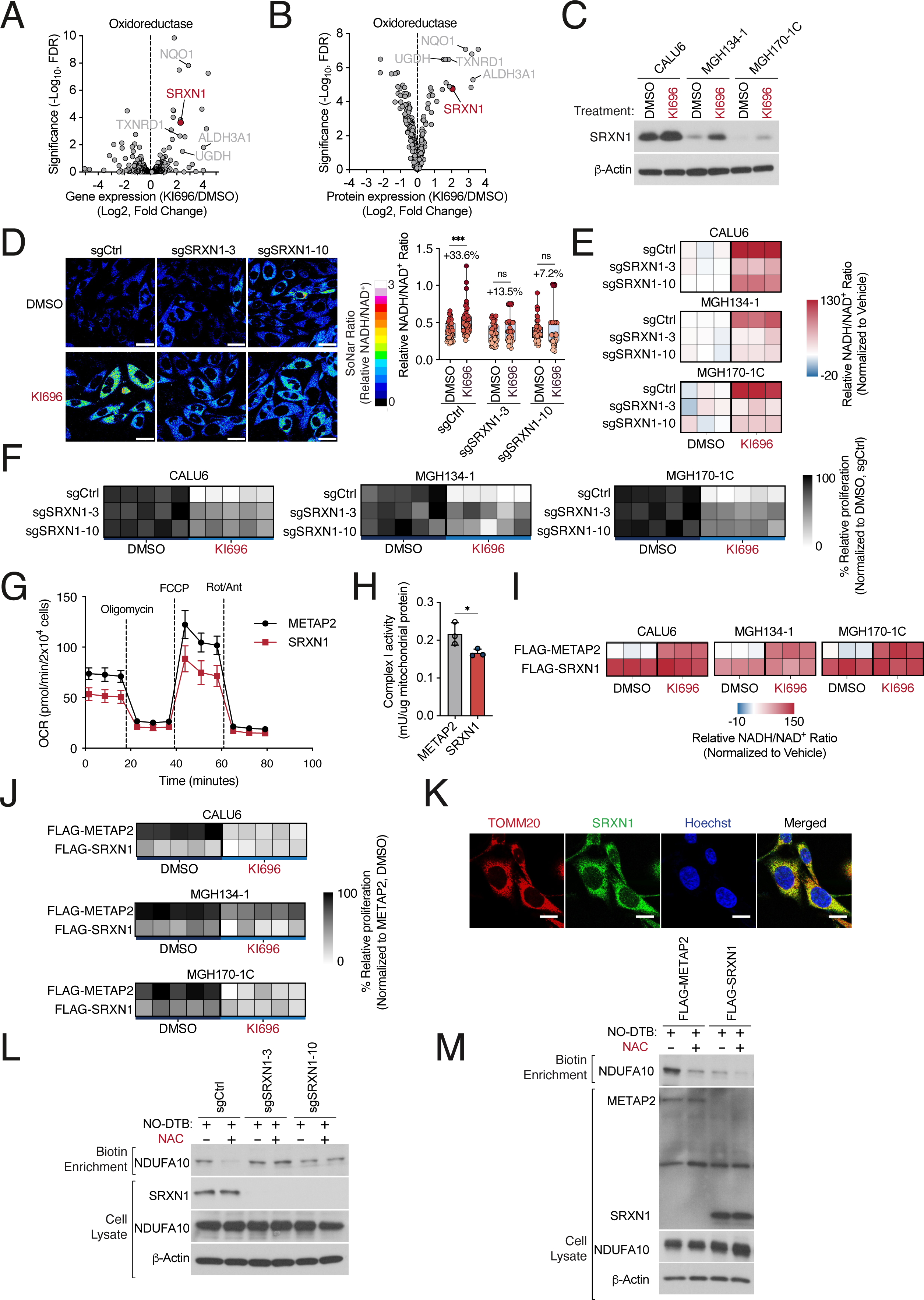
SRXN1 promotes regulates NDUFA10 oxidation. (**A, B**) SRXN1 is a NRF2-regulated oxidoreductase. Gene (**A**) and protein expression (**B**) of NRF2-regulated oxidoreductases following NRF2 activation in CALU6 cells. CALU6 cells were treated with 1 µM KI696 for 48 hrs and gene and protein expression were determined by RNAseq and total proteomics, respectively (adapted from Weiss-Sadan et al. (*77*)). (**C**) SRXN1 expression is induced by NRF2 activation in NSCLC cells. The indicated cell lines were treated with 1 µM of KI696 for 48 hrs and SRXN1 levels were determined by immunoblot. (**D-F**) Depletion of SRXN1 mediates resistance to NRF2 induced NADH-reductive stress in NSCLC cell lines. Cells stably expressing the indicated sgRNAs targeting SRNX1 or a control were treated with 1 µM of KI696 for 48 hrs and the NADH/NAD^+^ ratio was determined by immunofluorescent imaging in CALU6 (**D**) or by flow cytometry (**E**) in cells expressing SoNar, by comparing the emission intensity of 405 nm (NADH binding) vs. 488 nm (NAD^+^ binding). Cellular proliferation (**F**) was determined 120 hrs post treatment by measuring relative DNA levels using Hoechst 33342. (**G**) SRXN1 expression suppresses mitochondrial respiration. Representative Seahorse traces of oxygen consumption rate (OCR) in cells expressing FLAG-METAP2 (control) or FLAG-SRXN1. (**H**) SRXN1 expression decreases Complex I activity. Complex I activity was measured in cells expressing FLAG-METAP2 or FLAG-SRXN1 and normalized to mitochondrial protein. (**I, J**) SRXN1 over-expression results in NADH-reductive stress and a proliferation defect. Cells expressing METAP2 (control) or SRXN1 along with SoNar were treated with 1 µM of KI696 for 48 hrs. The NADH/NAD^+^ ratio determined by flow cytometry analysis (**I**), and cellular proliferation assays (**J**) following SRXN1 expression were performed as described in (**F**), respectively. (**K**) Immunofluorescence images of CALU6 cells stained with antibodies for TOMM20 and SRXN1. (**L, M**) NAC enhances SRXN1 activity. Measurement of NO-DTB labeling of NDUFA10 in CALU6 cells expressing the indicated sgRNAs following NRF2 treatment with 1 µM KI696 for 48 hrs followed by 10 mM NAC for 1 hr (**L**). Cellular lysates from CALU6 cells were treated with 500 µM NO-DTB and NDUFA10 reduction was determined by immunoblot following streptavidin-enrichment. CALU6 cells expressing FLAG-METAP2 or FLAG-SRXN1 were treated with NAC, sulfinylated proteins were enriched with the NO-DTB probe, and NDUFA10 was detected by immunoblotting (**M**). Statistical significance was determined by Student’s *t* test (two-tailed, unpaired). ns, not significant; * *p* < 0.05; *** *p* < 0.001. Scale bar: 25 µm (**D**) or 10 µm (**K**).

**Fig. S8:**
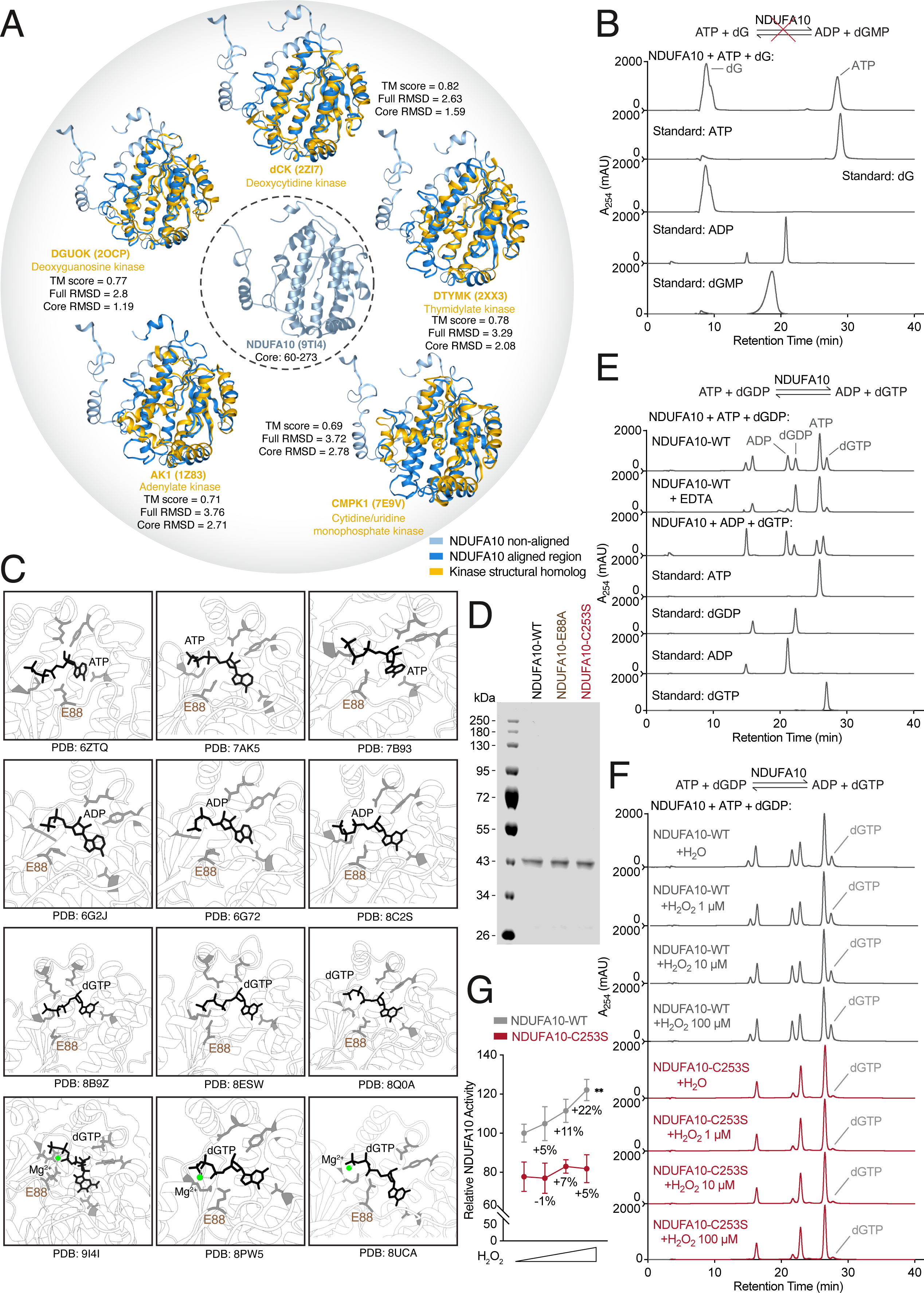
NDUFA10 catalyzes dGTP production in an oxidation-dependent manner. (**A**) Structural comparison of NDUFA10 with representative deoxynucleoside and nucleotide kinases. NDUFA10 is shown in blue (dark blue: aligned region, light blue: non-aligned region) and structural-related nucleotide/nucleoside kinases in yellow. Template Modeling score (TM-score), full root-mean-square deviation (RMSD), and core RMSD are indicated for each comparison. (**B**) NDUFA10 does not exhibit detectable dGK activity. Representative chromatograms (n=3) from reactions containing 1.5 µM NDUFA10, 1 mM ATP, 1 mM dG, and 10 mM MgCl_2_, compared with 1mM ATP, dG, ADP, and dGMP standards (see also Methods). (**C**) Representative ligand-bound NDUFA10 conformations observed in published mammalian Complex I structures. ATP-, ADP-, and dGTP-bound states from the indicated PDB entries are shown, highlighting the NDUFA10 binding pocket and the positioning of E88 and Mg²⁺. (**D**) Coomassie staining of recombinant NDUFA10 and associated variants (resi: 36-355) purified from E. Coli. (**E**) NDUFA10 catalyzes conversion of ATP and dGDP to dGTP in a Mg²⁺-dependent manner. Representative chromatograms (n=3) from reactions containing 1.5 µM NDUFA10 in the presence or absence of 10 mM EDTA, alongside ATP, dGDP, ADP, and dGTP standards. (**F, G**) Oxidation preserves NDUFA10 catalytic activity. Representative chromatograms (n=3) showing dGTP production by NDUFA10 WT or C253S following the indicated H_2_O_2_ treatments for 30 min (**F**), and quantification of the impact of H_2_O_2_ on kinase activity (**G**). Statistical significance was determined by Student’s *t* test (two-tailed, unpaired). ** *p* < 0.01.

**Fig. S9:**
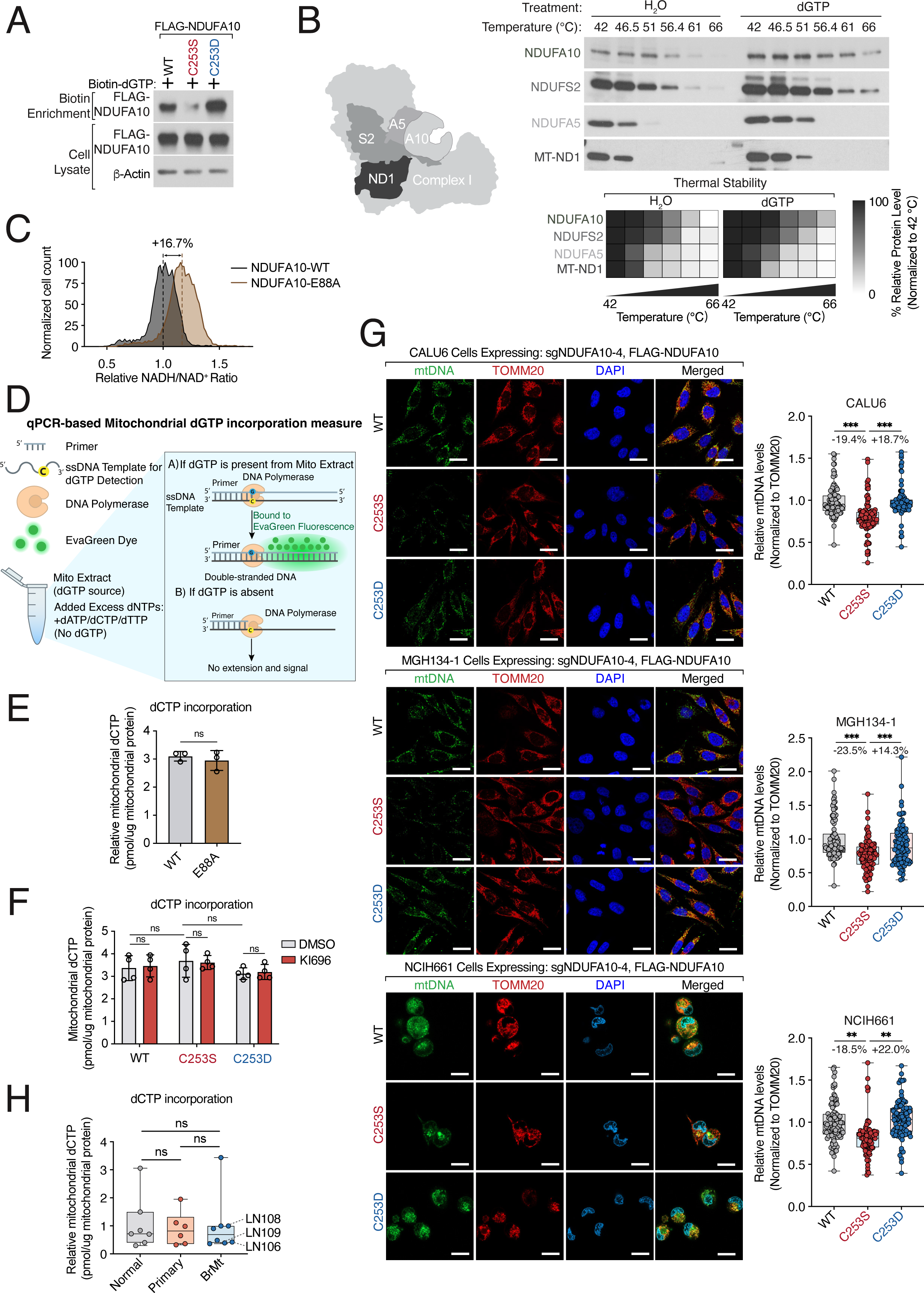
NDUFA10 oxidation supports mtDNA maintenance. (**A**) NDUFA10 cysteine oxidation regulates dGTP binding. CALU6 cells expressing different FLAG-NDUFA10 oxidation mimitec were lysed and the interaction between biotinylated dGTP and FLAG-NDUFA10 was determined by immunoblot analysis following streptavidin enrichment. (**B**) dGTP binding alters NDUFA10 conformation. Left, schematic of NDUFA10 interacting partners. Middle, thermal stability of NDUFA10 following in lysate treatment with 100 µM dGTP for 30 min was determined by incubation at the indicated temperatures, centrifugation and immunoblot analysis of the indicated proteins. Bottom, heatmap depicting relative abundance of Complex I proteins at each temperature. (**C**) NDUFA10-kinase dead mutant increases NADH levels. The NADH/NAD^+^ ratio was measured in CALU6 cells co-expressing PAM-mutants of NDUFA10 variants along with sgNDUFA10 and the SONAR reporter. (**D**) Schematic of a qPCR-based assay to measure incorporation of dGTP into mtDNA adapted from Purhonen et al. (*108*). Mitochondrial extracts were prepared, and dGTP was quantified by DNA polymerase-dependent primer extension measured by EvaGreen fluorescence. (**E**) NDUFA10 does not alter dCTP levels in mtDNA. Relative mitochondrial dCTP was measured in cells expressing NDUFA10 WT or Mg²⁺-binding mutant E88A as described in (**D**). (**F**) Lowering ROS level does not alter mitochondrial dCTP across the indicated NDUFA10 variants. Relative mitochondrial dCTP was measured in cells expressing WT, C253S, or C253D NDUFA10 in the background of endogenous NDUFA10 depletion following treatment with 1 µM of KI696 for 48 hrs. (**G**) Oxidation state of NDUFA10 Cys253 regulates mtDNA levels. Representative images of mtDNA, TOMM20, and DAPI staining in CALU6, MGH134-1, and NCIH661 cells expressing sgNDUFA10 together with PAM mutant version of WT, C253S, or C253D NDUFA10. Quantification of relative mtDNA volume is shown at right. (**H**) Brain metastases do not exhibit increased mitochondrial dCTP relative to primary tumors. dCTP incorporation into mtDNA was measured as described in (**D**). Statistical significance was determined by Student’s *t* test (two-tailed, unpaired). ** *p* < 0.01; *** *p* < 0.001. Scale bar: 25 µm.

**Fig. S10:**
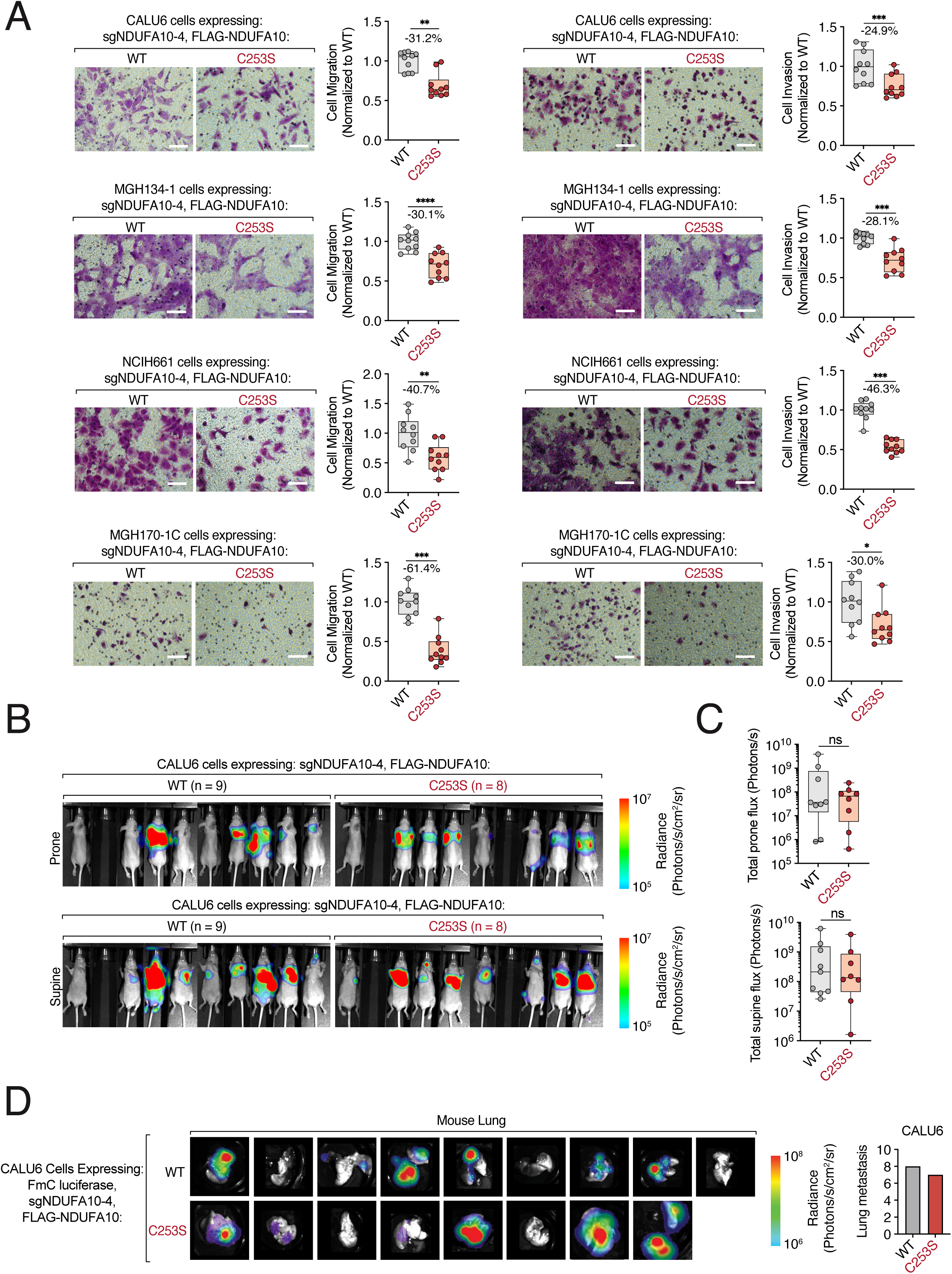
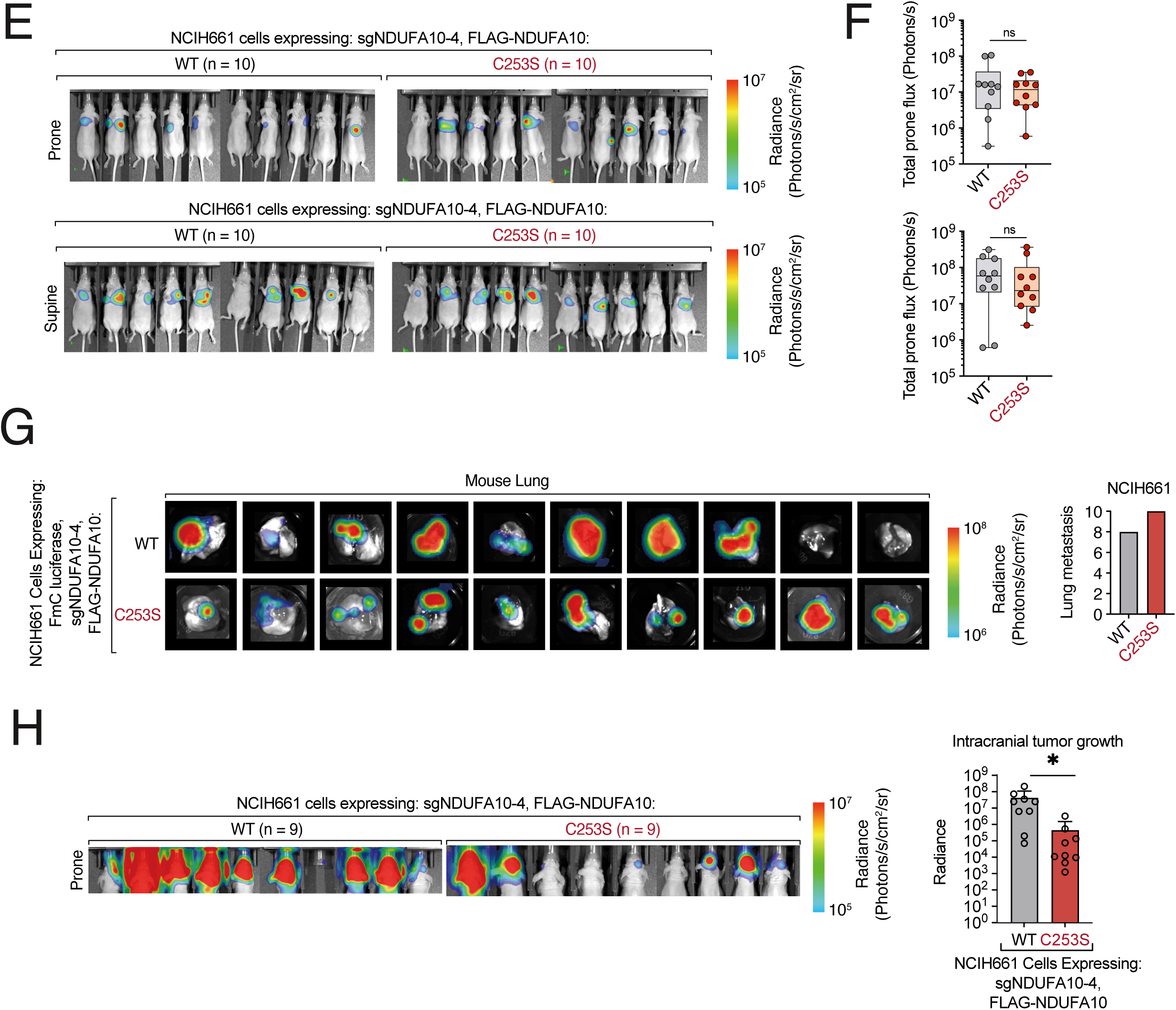
NDUFA10•C253 oxidation supports brain metastasis. (**A**) NDUFA10 oxidation facilitates *in vitro* NSCLC cell migration and invasion. The indicated NSCLC cells were depleted of endogenous NDUFA10 and transduced with a PAM-mutant of FLAG-NDUFA10 redox variants (WT and C253S) followed by analysis of cell migration and Matrigel invasion. Left, representative images of cell migration and invasion. Right, quantification of migration and invasion (see also Methods). (**B, C**) Whole-animal metastatic burden is not significantly changed in the CALU6 intracardiac model. *In vivo* bioluminescence images of whole body (**B**) and quantification (**C**) of mice intracardially injected with CALU6 cells depleted of endogenous NDUFA10. (**D**) NDUFA10 oxidation does not alter lung colonization in the CALU6 intracardiac model. Left, mouse lungs were isolated from mice treated as described in (**B**) and *in vitro* bioluminescence was determined. Right, quantification of intracardiac tumor growth in lung. (**E**) Whole-animal metastatic burden is not significantly changed in the H661 intracardiac model. Representative bioluminescence images of mice following intracardiac injection of NCIH661 cells expressing sgNDUFA10 together with FLAG-tagged WT or C253S NDUFA10. (**F**) Quantification of whole-animal bioluminescence from mice shown in (**E**). (**G**) NDUFA10 reduction does not alter lung colonization in the NCIH661 intracardiac model. Left, representative *ex vivo* bioluminescence images of lungs from mice injected intracardially with NCIH661 cells expressing sgNDUFA10 together with FLAG-tagged WT or C253S NDUFA10. Right, quantification of lung metastatic burden is shown at right. (**H**) NDUFA10 reduction suppresses intracranial tumor growth in NCIH661 models. Left, representative mouse image from intracranial implantation experiments using NCIH661 cells expressing sgNDUFA10 together with FLAG-tagged WT or C253S NDUFA10 PAM mutants. Right, quantification of intracranial tumor growth by bioluminescence imaging. Statistical significance was determined by Student’s *t* test (two-tailed, unpaired), or Fisher’s exact test. * *p* < 0.05; ** *p* < 0.01; *** *p* < 0.001. Scale bar: 100 µm.

## Materials and Methods

### Tissue culture

All cells were maintained at 37°C with 5% CO_2_. HEK-293T, LN-001, LN-084, and LN-088 cells were grown in DMEM (Corning) supplemented with 10% fetal bovine serum (FBS, Corning), Penicillin-Streptomycin (100 mg/ml, Millipore) and L-glutamine (2 mM, Corning). All other cell lines were grown in RPMI-1640 (Corning) media containing 10% FBS (Corning), Penicillin-Streptomycin (100 mg/ml, Millipore) and 1% GlutaMax (Gibco). All cell lines were routinely tested for Mycoplasma and if not noted elsewhere were obtained from American Tissue Type Collection (ATCC). All *MGH* cell lines were kindly provided by Dr. Aaron N. Hata. Whenever thawed, cells were passaged at least three times before being used in experiments.

### cDNA cloning and mutagenesis

cDNAs were amplified using Q5 High-Fidelity 2X master mix (NEB) and subcloned into pLJM1 (Addgene) or pLentiCRISPRv2 (Addgene) by T4 ligation or Gibson cloning (NEB). Site directed mutants were generated using QuikChange XLII site-directed mutagenesis (Agilent) or Gibson cloning (NEB), using primers containing the desired mutations. The sequences of primers used in this study can be found in Supplementary Table S7. The SoNar cDNA is previously described (*72*). All constructs were verified by DNA sequencing.

### Lentivirus production

Lentivirus was produced as previously described (*53*). Briefly, mammalian lentiviral particles harboring sgRNA-encoding plasmids or cDNA-encoding plasmids were co-transfected with the psPAX2 envelope and VSV-G packaging plasmids into actively growing HEK-293T cells using Xtremegene-HP (Sigma-Aldrich) transfection reagent as previously described (*109*). Virus-containing supernatants were collected 48 hrs after transfection and filtered to eliminate cells. Target cells were infected in the presence of 10 μg/ml polybrene (Millipore) at a concentration of 2.5 x 10^5^ cells/well. 24 hrs post-infection, fresh media was added to the infected cells which were allowed to recover for an additional 24 hrs. Cells were selected with puromycin (Sigma-Aldrich), blasticidin (Sigma-Aldrich), or Hygromycin B (Sigma-Aldrich) and analyzed 3-10 days after selection was initiated. The sequences of sgRNAs used in this study can be found in Supplementary Table S7.

### Patient samples

Lung cancer brain metastasis tissue samples were collected in accordance with ethical guidelines from the Declaration of Helsinki and Belmont Report and under the Tissue Bank for Neurological Disorders protocol (#10-417), which is approved by the Dana-Farber Cancer Institute’s Institutional Review Board. 19 patients who were undergoing craniotomy for suspected or confirmed lung cancer brain metastases at Massachusetts General Hospital (MGH), and who were able and willing to provide written, informed consent, were prospectively enrolled into the study. Immediately following craniotomy for tumor resection, lung cancer brain metastasis tissue in excess for clinical care was approved for research use by Board-certified pathologists at MGH, where it was immediately transported to the laboratory and snap frozen using an isopropyl alcohol and dry ice slurry. Tissue was stored at -80° C until further analysis. Primary lung tumor and matched adjacent normal lung tissue samples were obtained through the MGH Lung Tumor Bank under protocol #13-416. A total of 62 snap-frozen samples, comprising 31 matched pairs of primary lung tumor and normal lung tissue, were approved for release. Each individual tumor or normal tissue sample was required to have a minimum weight of 50 mg to be considered usable for this project. De-identified clinical data was collected from patient charts, and pathology reports were reviewed in detail to ascertain histological diagnosis, clinical features and molecular and mutational analysis performed as part of clinical care.

### Cryogenic milling

Frozen tissues were maintained on dry ice or in liquid nitrogen throughout processing to minimize thawing and artifactual oxidation. Cryo-mill tubes, stainless steel grinding beads, tube holders, and metal grinding surfaces were pre-chilled in liquid nitrogen before use. Frozen tissue was transferred into a pre-chilled plastic pouch, sealed, and manually pulverized on a liquid nitrogen-cooled metal surface using a pre-chilled hammer. The resulting frozen tissue powder was transferred through a pre-chilled funnel into cryo-mill tubes containing two 5-mm stainless steel grinding beads. Samples were immediately processed by cryogenic milling using a CryoMill cryogenic grinder (Retsch). Cryo-milling was performed using program P4 with 3 cryo-cycles, a starting frequency of 5 Hz increasing to 20 Hz, and an 11-min milling time. After milling, tubes were briefly vented to release pressure, returned immediately to dry ice, and stored at −80°C until lysis and downstream analysis.

### iso-TMT sample preparation

Iso-TMT samples were prepared as previously described(*58, 110, 111*). In brief, two independent biological replicates of each NSCLC cell lines (Supplementary Table S1) were treated in independent with 1 µM KI696 (MedChem Express) or 5 µM Doxycycline Hydrochloride (DOXY) (Sigma-Aldrich) for 48 hours, or treated with 10 mM N-acetyl-L-Cysteine (NAC) (Sigma-Aldrich), 10 mM Glutathione reduced ethyl ester (GSHee) (Sigma-Aldrich), 250 µM Trolox (Cayman) for 1 hr, or 200 µM H_2_O_2_ for 30 min. Cells were washed once with ice-cold PBS, snap-frozen in liquid nitrogen, and stored at -80 °C until use. Frozen cell pellets or cryomilled tissues were lysed in DPBS supplemented with Benzonase (Santacruz) and protease inhibitors (Roche) using a chilled bath sonicator (Q700, Qsonica) and centrifuged for 3 min at 300 x g. Proteins were quantified by BCA assay (Thermo Fisher Scientific) and a total of 50 µg protein extracts were used for each TMT channel. Lysates were treated with 1 mM DBIA treatment for 1 hr or for unenriched analysis (e.g., total proteome analysis) samples were directly alkylated and enriched with SP3 as described below. Following DBIA incubation, lysates were reduced with 5 mM 5-tris(2-carboxyethyl) phosphine hydrochloride (TCEP) (Sigma-Aldrich) for 2 min at room temperature, followed by alkylation using 20 mM chloroacetamide (Sigma-Aldrich) for 30 min in the dark at room temperature. SP3 magnetic beads (Cytiva) were prewashed with LC-MS grade water and 250 µg combined SP3 beads (1:1 hydrophobic:hydrophilic) and LC-MS grade ethanol were added to each sample to reach a final concentration of 50% ethanol(*112*). SP3 protein binding occurred for 30 min at room temperature and beads were subsequently washed 3 times with 80% ethanol and resuspended with 175 µl of Trypsin/Lys-C (1 µg, Thermo Fisher Scientific) in 200 mM EPPS (pH 8.4)/5 mM CaCl_2_. For total proteomic analysis, peptides were dried as described below. Proteins were digested overnight (16 hrs) at 37 °C and digested peptides were enriched with streptavidin magnetic beads (Cytiva) for 1 hr at room temperature. Beads were subsequently washed three times with DPBS, twice with HPLC grade-water (Sigma-Aldrich). Peptides were eluted with 50 % acetonitrile (Sigma Aldrich), 0.1% formic acid (Thermo Fisher Scientific), and dried using a SpeedVac (Thermo Fisher Scientific). Cysteine-enriched peptides were reconstituted with 30 % acetonitrile, 70 % 200 mM EPPS pH 8.4 and labeled with 25 µg of TMT reagent (Thermo Fisher Scientific) per channel for 75 min at room temperature with rotation. Labeling was terminated by the addition of 5% hydroxylamine (Acros Organics) for 15 min followed by addition of 10% formic acid. Peptides were labeled with TMTpro16-plex reagents (Thermo Fisher Scientific), pooled, and dried using a SpeedVac (Thermo Fisher Scientific). Peptides were desalted with stage tips using the following procedure: peptides were reconstituted with 5% acetonitrile/0.1 % formic acid and loaded onto C18 Micro Spin columns (Nest Group) pre-equilibrated with LC-MS grade methanol (Fisher Chemical) and LC/MS-grade water containing 0.1% formic acid. C18 spin columns were washed 10 times with LC/MS grade water containing 0.1 % formic acid, subsequently eluted with 80% acetonitrile and 0.1% formic acid, and dried using a SpeedVac (Thermo Fisher Scientific).

### MS data acquisition and processing

All the mass spectrometry samples were analyzed as described previously (*53*) on an Orbitrap Eclipse^TM^ Tribrid^TM^ Mass Spectrometer in-line with an Easy NanoLC-1200 system (Thermo Fisher Scientific)(*110*). Peptides were separated on a 75-µm capillary column packed with 50 cm of C18 resin (2 µm, 100 Å; Thermo Fisher Scientific) using a 180 min gradient of 4-35% acetonitrile in 0.1% FA with a flow rate of 300 nL/min. Eluted peptides were acquired by data-dependent acquisition and quantified using the synchronous precursor selection (DDA-SPS-MS3) method for TMT quantification. Briefly, MS1 spectra were acquired in the scan range of 400-1400 m/z at an orbitrap resolution of 120,000 with 50 ms as a maximum injection time with high-field asymmetric-waveform ion-mobility spectrometry (FAIMS) values at -40, -50, and -70 compensation voltage (CV). MS2 spectra were acquired by the selection of the top twenty most abundant features via collisional induced dissociation in the ion trap using a normalized automatic gain control (AGC) target of 100, normalized collision energy, quadrupole isolation width of 0.7 m/z and a maximum ion accumulation time of 50 ms. Next, SPS-MS3 scans were performed using 20 b- and y-type fragment ions as precursors with a normalized HCD collision energy of 55 and AGC target of 500 having the maximum injection time as 250 ms with 1.2 isolation window.

Acquired RAW data were analyzed using Proteome Discoverer v2.5 (Thermo Fisher Scientific) using the SequestHT module. Data was search against the UniProtKB human universal database (release_20210506) combined with the common Repository of Adventitious Proteins (cRAP, classes 1,2,3, and 5). Searches included both the canonical and isoform-inclusive reference proteomes; however, all downstream analyses were performed using the canonical reference proteome. Parameters were set as follows: MS1 tolerance of 10 ppm, MS/MS mass tolerance of 0.6 Da, trypsin (full) digestion with a maximum of zero missed cleavages, minimum peptide length of 6 amino acids, and maximum of 144 amino acids. Cysteine carbamidomethylation (57.021 Da) and methionine oxidation (15.995 Da) were set as dynamic modifications. Lysine- and N-terminus-TMTpro modification (304.207 Da) were set as static modification. A false discovery rate (FDR) of 1% was set for peptide-to-spectrum matches using the Percolator algorithm and for protein assignment. Reporter ion quantification was based on signal over noise (S/N) with the co-Isolation Threshold at 50, S/N threshold at 10, and SPS mass matches threshold at 50%. For the TMT reporter ion quantification, all the identified peptide spectral matches (PSMs) from the MS3 scans were extracted by an in-house program and the reporter ion intensities were adjusted for impurity correction according to the manufacturer’s specifications. For quantification of each MS3 spectrum, a total sum signal-to-noise of all reporter ions of 100 (TMT16-plex) was used. Abundances were normalized to total peptides.

### Genome-wide CRISPR screen

Genome-wide CRISPR screens were conducted as previously described(*113, 114*). Briefly, CALU6 cells were infected with a genome-wide CRISPR knockout pooled library (Brunello)(*115*)(Addgene), ensuring a multiplicity of infection ∼0.3 following 3-4 days puromycin selection. Cells were allowed to recover for 2-3 days, and an initial input was taken with the number of infected cells corresponding to >1000 x the size of the library. CRISPR library infected cells were split every other day to a defined concentration of 5 x 10^5^ cells per ml to maintain ideal cell concentration and growth. The screen was initiated by treating cells (∼80 x 10^6^) with 1 µM KI696, 2.5 µM DOXY, 10 mM NAC, 5 mM GSHee, or 100 µM Trolox, and this cell number and compound concentration were maintained for 11 population doublings. At the end of the screen, cells were harvested, and genomic DNA was extracted using Macherey Nagel Blood XL kit (Macherey). Libraries were generated from each sample by PCR based amplification of the sgRNA amplicon from 200 µg of genomic DNA using custom PCR primers harboring an index primer and Illumina 5’ and 3’ adaptors(*116*). Libraries were pooled and analyzed on a NextSeq500 (Illumina) using single end 75 bp reads. sgRNAs were mapped and quantified using the Screen Processing pipeline. To calculate the CRISPR score for each sgRNA, log2 normalized vehicle and compound sgRNA values were subtracted from corresponding input values and the median sgRNA value for each gene was determined.

### H_2_DCFDA measurement

H_2_DCFDA (Thermo) analysis was conducted as previously described(*48*). CALU6, MGH134, and NCIH661 cells were treated as described in the text and washed with prewarmed PBS and harvested by centrifugation at 1200 *g* at room temperature for 2 mins. The cell pellet was resuspended in PBS with 1 µM of CM-H_2_DCFDA and incubated for 45 min in a 37°C incubator with controlled CO_2_ levels (5%). Subsequently cells were washed with PBS. Changes in CM-H_2_DCFDA fluorescence were determined by flow cytometry using Aurora (Cytek). Data was analyzed using Flowjo v10.6 for FITC intensity.

### Cell proliferation assays

Unless otherwise noted, cells were pre-treated for 48 hrs in 6-well plates, prior to the onset of proliferation assays. At the onset of a proliferation assay, cells were cultured in 96-well plates at 2.5 x 10^3^ cells per well in 100 µl of medium and the indicated compound or vehicle control was added for compounds where pre-treatment was required. For all other compounds, agents were administered 24 hrs after cell seeding. To quantify cell proliferation Hoechst (Sigma-Aldrich) was added to each well of samples after 120 hrs of treatment and fluorescence was determined on SpectraMax M5 plate reader according to the manufacturer’s directions (Molecular Devices).

### Seahorse flux analyses

Mitochondrial oxygen consumption rates were measured using an XFe96 Extracellular Flux Analyzer (Agilent) as previously described (*77*). Cells were seeded on poly-L-lysine-coated Seahorse XF96 cell culture plates and treated with the indicated compounds as described in the text. For KI696 and DOXY conditions, cells were treated for 48 h before analysis. Before measurement, culture medium was replaced with bicarbonate-free RPMI supplemented with 2 mM L-glutamine and 10 mM D-glucose, and cells were equilibrated in a non-CO₂ incubator. Oxygen consumption rates were measured at baseline and following sequential injection of oligomycin, FCCP with sodium pyruvate, and antimycin A/rotenone. Basal respiration, ATP-linked respiration, and maximal respiration were calculated from oxygen consumption traces and normalized to total protein abundance measured by BCA assay.

### Mitochondria isolation

Cell pellets were gently re-suspended and incubated on ice as described in (*77*), then transferred to a ball-bearing cell homogenizer (Active Motif) and homogenized on ice. To confirm lysis efficiency, 10 μl of cell lysate was stained with 0.5% trypan blue and viewed under a light microscope. Homogenization was continued until 95% cell membrane breakage was achieved (typically 10–15 passages through the homogenizer). The homogenate was centrifuged at 600 x *g* for 10 min at 4 °C to remove nuclei and membranous debris, and the supernatant containing mitochondria was collected, and the density gradient ultracentrifugation was conducted as previously described (*117*). Briefly, five gradient solutions were prepared by mixing gradient dilution buffer with the OptiPrep medium. The diluted OptiPrep density gradient solutions were carefully overlaid in descending concentration order in a 7 ml ultracentrifuge tube (Beckman). Next, the prepared supernatant was overlaid on top of the density gradient. The tube was centrifuged at 145,000 x *g* for 4 hrs at 4 °C. A total of 10 fractions of 600 μl each were carefully withdrawn from the tops of the gradients using extra-long micropipette tips (Thermo Fisher Scientific). Each isolated fraction was mixed with 1000 μl PBS and centrifuged at 20,000 x *g* for 30 min at 4 °C to collect the pellets as mitochondrial fractions and stored at −80°C.

### Complex I activity assays

Complex I activity assays were performed as previously described (*118*). In brief, mitochondrial pellets were resuspended in 0.5 ml of 10 mM ice-cold hypotonic Tris buffer (pH 7.6). A 10 μl aliquot was taken for total protein concentration measurements according to the Bradford (Bio-Rad) method. The mitochondrial solution was thawed and subjected to three cycles of freeze-thawing in liquid nitrogen to disrupt the mitochondrial membranes. 20-50 μg of isolated mitochondrial protein was added to 700 μl of distilled water in a 1 ml cuvette. Subsequently 100 μl of potassium phosphate buffer (0.5 M, pH 7.5), 60 μl BSA (50 mg/ml), 30 μl of KCN (10 mM) and 10 μl of NADH (10 mM) were added, and the volume was adjusted to 994 μl with distilled water. A separate cuvette containing the same quantity of reagents and sample but with the addition of 10 μl of 1 mM rotenone solution was prepared in parallel. The cuvettes were mixed by inverting with Parafilm and the baseline was read at 340 nm for 2 min. The reaction was started by adding 6 μl of ubiquinone_1_ (10 mM), mixed by inverting the cuvette using Parafilm, and followed by the decrease of absorbance at 340 nm for 2 min. Changes in complex I activity were analyzed using Prism v10 (GraphPad) software.

### Confocal imaging of cell lines expressing SoNar reporter

NADH/NAD^+^ levels were measured as previously described (*48, 77*). NSCLC cell lines expressing the indicated SoNar reporter was seeded on poly-lysine coated 8-well chamber (iBidi) at 2 x 10^4^ cells per well and treated with compounds as described in the text. For KI696 and DOXY treatments, cells were pre-treated for 2 days with 1 µM KI696 or 5 µM DOXY prior to seeding on glass bottom dishes. Dishes were firmly mounted the stage adaptor of Zeiss 710 Laser Scanning Confocal microscope (Carl Zeiss Inc.). Constant temperature (37°C), humidity, and 5% CO_2_ atmosphere are maintained throughout the duration of cell imaging. Images were acquired using a 63X oil objective. For NADH/NAD^+^ ratio determination, relative NAD^+^ levels were measured by exciting SoNar expressing cells with a 488 nm laser and measuring emission at 500-520 nm range. Relative NADH levels was determined by exciting SoNar expressing cells with a 405 nm laser and measuring emission at 500-545 nm range. Acquisition parameters were kept identical between samples. Ratiometric images of SoNar were processed using ImageJ (NIH, v2.0.3). Threshold images after subtraction of background were split into two different channels, divided with Image Calculator. 32-bit ratiometric images were generated and presented in 16 colors mode from the Lookup Tables.

### Flow cytometry analysis

NADH/NAD^+^ ratio was determined by flow cytometry as previously described (*77*). NSCLC cells expressing the indicated SoNar reporters were seeded at 0.25 x 10^6^ cells/well in a 6-well plate for 48 hrs. For KI696 of DOXY treatments, cells were pre-treated for 2 days with 1 µM KI696 or vehicle control. Cells were dislodged by trypsin digestion, resuspended in FACS buffer (PBS with 1% FBS and 2 mM EDTA). The fluorescent signal at 530 nm following excitation at 405 nm (NADH binding) or 488 nm (NAD^+^ binding) was measured by an Aurora (Cytek) flow cytometer. The ratio of λex = 488 nm/ λem = 530 nm to λex = 405 nm/ λem = 530 nm signal was determined using Flowjo v10.6.

### Immunofluorescence

Immunofluorescence was performed as described previously (*48*). In brief, CALU6 cells expressing the indicated proteins were seeded on poly-lysine coated 8-well chamber (iBidi) at 2 x 10^4^ cells per well and treated with compounds as described in the text. For KI696 treatments, cells were pre-treated for 2 days with 1 µM KI696 or vehicle control prior to seeding on glass bottom dishes. Cells were fixed with 4% PFA (EMS) for 15 min and permeabilized with 0.1% Triton X-100 in PBS on ice for 10 min. The slides were rinsed with PBS and incubated with primary antibodies in 4% BSA for 1 hr. Following three PBS washes, the slides were incubated with secondary antibodies conjugated to the Alexa Fluor 488 and 647 fluorophores (Invitrogen) for 30 min at room temperature. The slides were rinsed and covered by PBS. Cells were imaged on a Zeiss LSM 710 laser scanning confocal microscope under 63x oil objective. Images were processed using ZEN 2.6 Image software (Carl Zeiss Inc.) and ImageJ (NIH).

### Cell lysis and immunoprecipitations

Cell lysis and immunoprecipitations were performed as described previously (*111*). In brief, the indicated cell lines were rinsed once with ice-cold PBS and lysed using a chilled bath sonicator (Q700, QSonica) in Triton lysis buffer (1% Triton X-100, 40 mM HEPES pH 7.4, 2.5 mM MgCl_2_) supplemented with protease inhibitors (Roche), 10 mM Sodium Fluoride (Sigma-Aldrich), 1 mM Sodium Orthovanadate (Sigma-Aldrich), and Benzonase (Santa Cruz). The soluble fractions of cell lysates were isolated by centrifugation at 13,000 rpm in a microcentrifuge for 10 min, normalized to 1 mg/ml. Proteins were denatured by the addition of 5X sample buffer and boiling for 5 min. For FLAG immunoprecipitations, anti-FLAG M2 resin (Sigma-Aldrich) was added to the pre-cleared lysates and incubated for 3 hrs at 4°C. Following immunoprecipitation, beads were washed once with Triton IP buffer followed by 3 times with Triton IP buffer supplemented with 500 mM NaCl. Loading buffer was added to the immunoprecipitated proteins which were subsequently denatured by boiling for 5 min. Samples were resolved by 10%–20% SDS-PAGE and analyzed by immunoblotting.

### Thermal stability assays

Thermal stability assays were performed as previously described (*53*). CALU6 cells stably expressing the indicated proteins following the indicated treatments as described in the text were harvested and the pellets lysed with DPBS supplemented with protease inhibitor (Roche), Sodium Fluoride (Sigma-Aldrich), and Sodium Orthovanadate (Sigma-Aldrich), using a chilled bath sonicator (Q700, QSonica). The lysates were clarified by centrifugation at 300 *g* for 3 min. The supernatants were diluted to 1.25 mg/mL using DPBS and aliquoted at 50 µL/well in a PCR strip. For thermal stability assays, some lysates were treated with 100 µM H_2_O_2_ or 100 µM dGTP at room temperature for 30 min as indicated in the text. The samples were then heated to the indicated temperature in a BioRad T100 Thermal Cycler (BioRad) for 3 min. Samples were cooled at room temperature for 3 min, incubated on ice for 3 min, transferred into 1.5 mL tubes, and centrifuged at 21,000 *g* for 1 hr at 4°C. The soluble fractions were collected for immunoblot analysis. The band intensity was assessed using ImageJ analysis and normalized to the value at the lowest temperature used for each experiment (i.e., 42 °C). The melting curves of each protein were generated by Prism v10 (GraphPad) software.

### Detection of NDUFA10 sulfinylation

Detection of NDUFA10 sulfinylation by NO-DTB labeling was performed as previously reported (*33*). In brief, the indicated NSCLC cells were treated with the indicated compounds as described in text. For mouse tissues, frozen tissue powder was generated by cryogenic milling as described above. Cell samples were washed with ice-cold PBS and lysed using a chilled bath sonicator (Q700, QSonica) in Triton IP buffer (1% Triton X-100, 5 mM MgCl2, 40 mM Hepes pH 7.4, 10 mM KCl) supplemented with protease inhibitors (Roche), phosphatase inhibitors (Roche), Benzonase (Santa Cruz), and 5 mM DTT. Lysates were clarified by centrifugation at 13,000 rpm for 10 min. Free thiols were trapped by incubation with 2 mM of 4,4’-dithiodipyridine (4-DPS) at room temperature for 1 h and subsequently buffer exchanged using one Micro Bio-Spin column pre-equilibrated with 100 mM HEPES, pH 8.5, 100 mM NaCl. DPS-free lysates were then reacted with 500 µM NO-DTB in the dark at room temperature with rotation for 1 hr. Total protein was purified by the addition of a chloroform-methanol solution (4:4:1, methanol, water, chloroform) to each sample and precipitated following centrifugation at 4200 rpm for 10 min. The protein disc was isolated, washed once in methanol, resuspended in Buffer X1 (9 M Urea, 10 mM DTT, 50 mM tetramethylammonium bicarbonate (TEAB)), and incubated for 20 min at 65°C. NO-DTB modified proteins were then enriched by the addition of streptavidin beads (Thermo Fisher Scientific) and following 2 hrs incubation, beads were washed twice with 0.1% IGEPAL (Sigma-Aldrich), PBS, and Triton IP buffer supplemented with 500 mM NaCl. 5 x loading dye was added to the samples. Proteins were then resolved by SDS-PAGE and analyzed by immunoblotting. NO-DTB enrichment was normalized to input NDUFA10 abundance for comparison across mouse tissues.

### DAAO-mediated localized H₂O₂ generation

Localized H₂O₂ generation was performed using a D-amino acid oxidase (DAAO)-based system as previously described (*61*). Cells stably expressing FLAG-DAAO-NDUFA10 construct was treated with 10 mM D-alanine to induce localized H₂O₂ production or 10 mM L-alanine as a negative control for 3 hrs. To determine the location of FLAG-DAAO-NDUFA10, cells were fixed and processed for immunofluorescence as described above. For thermal stability analysis, cells were harvested after alanine treatment, lysed in DPBS supplemented with protease and phosphatase inhibitors, and subjected to thermal stability assays as described above. Protein stability was determined by immunoblotting and quantified using ImageJ.

### Mitochondrial dGTP incorporation and mtDNA quantification

Mitochondrial dGTP incorporation was measured using a modified qPCR-based primer extension assay as previously described (*108*). Mitochondrial extracts were prepared from the indicated cells or tissue specimens as described above. Mitochondrial dGTP was quantified by DNA polymerase-dependent primer extension using EvaGreen fluorescence, with relative signal normalized to the indicated control condition. Mitochondrial dCTP incorporation was measured in parallel as a control where indicated. For mtDNA quantification, total DNA was isolated from cells or tissues, and mitochondrial DNA abundance was measured by qPCR using primers targeting MT-ND1 and MT-CO1. Mitochondrial DNA levels were normalized to nuclear DNA using B2M and GAPDH as nuclear encoded controls.

### Structural analysis and visualization

Structural similarity between NDUFA10 and representative deoxynucleoside or nucleotide kinases was assessed using Foldseek (*119*). The NDUFA10 structure from the human Complex I cryo-EM structure PDB: 9TI4 was used as the query. Structural similarity was evaluated using the Template Modeling score (TM-score) and root-mean-square deviation (RMSD) reported by Foldseek, with core RMSD calculated in PyMOL after alignment of the conserved structural core. Mammalian Complex I structures were inspected for ligands bound within the NDUFA10 nucleotide-binding region (Fig. 6A, fig. S8C) (*87, 89, 90, 120–130*). Ligand-binding residues and distances to bound nucleotides in the indicated PDB structures were analyzed and structural scenes were generated using Chimera 1.17.3.1 (*131*).

### Recombinant NDUFA10 purification

Recombinant NDUFA10 purification was performed as described previously (*132*). Briefly, wild-type and mutant constructs cloned into the pET28a(+) plasmid were transformed into T7 expression-compatible competent *E. coli* cells (New England Biolabs) for recombinant protein production. Single colonies were inoculated into Luria⍰Bertani (LB) medium containing kanamycin (100 µg/mL) and grown overnight at 37 °C. Overnight cultures were diluted 1:100 into fresh LB medium containing 100 µg/mL kanamycin and grown at 37 °C until the optical density at 600 nm (OD600) reached 0.6. Protein expression was induced with 0.8 mM isopropyl-β-D-thiogalactoside (IPTG; Research Products International) at 16 °C for 12–16 h. Cells were harvested by centrifugation at 11,000 x g for 5 min at 4 °C, and the supernatant was discarded. The cell pellets were resuspended in 1X phosphate⍰buffered saline (PBS) and sonicated on ice using an ultrasonicator (Branson SFX550 Sonifier) at 38% power, with 2 cycles of 10 sec on/10 sec off. After centrifugation at 17,000 x g for 30 min at 4 °C, the supernatant was discarded. The pellets were resuspended in lysis buffer (40 mM imidazole, 20 mM sodium phosphate, 500 mM NaCl, 10 mM MgCl_2_, 10% Glycerol, and 6 M urea, pH 7.4). The lysate was incubated at room temperature for 45 min on a rotary shaker, followed by centrifugation at 17,000 x g for 30 min at 4 °C. The filtered supernatant was pooled, and equilibrated cobalt agarose beads (Thermo Fisher Scientific) were added and incubated at 4 °C for 45 min on a rotary shaker. Following five washes with wash buffer (40 mM imidazole, 20 mM sodium phosphate, 500 mM NaCl, 10 mM MgCl_2_, 10% Glycerol, and 6 M urea, pH 7.4), the proteins were eluted with elution buffer (450 mM imidazole, 20 mM sodium phosphate, 500 mM NaCl, 10 mM MgCl_2_, 10% Glycerol, and 6 M urea, pH 7.4) at 4 °C. The buffer was exchanged by dialysis (7,000 MWCO, Thermo Fisher Scientific), and urea and imidazole were removed gradually. The purity of the proteins was determined using 10% SDS⍰PAGE with Coomassie Blue staining.

### Recombinant NDUFA10 *in vitro* kinase assay

1.5 μM of purified NDUFA10 WT or mutant proteins were incubated with 1mM ATP and 1mM dG or 1mM dGDP in 20 mM sodium phosphate, 100 mM NaCl, 10 mM MgCl_2_, and 40 mM KCl (pH 7.4) at 37 °C for 30 min. The reaction was quenched by adding an equal volume of ice-cold methanol. The mixture was vortexed and then centrifuged at 21,130 × g for 20 min at 4 °C. The aqueous phase was collected and injected onto an ion-exchange column (SOURCE™ 15Q, Cytiva) at a flow rate of 0.4 mL min⁻¹ using an HPLC system. The column was equilibrated with 2 mM ammonium formate before HPLC analysis. The sample was eluted using a linear gradient from 2 to 600 mM ammonium formate over 25 min (*133*). Nucleotides were detected by UV absorbance at 254 nm using an Agilent 1100 HPLC instrument.

### Migration and Matrigel invasion assays

Migration and invasion assays were performed as previously described (*134*). Following construction of cell lines (fig. S10A), cells (1 x10^5^/well) were suspended in serum-free RPMI were seeded onto the upper chamber of 24-well transwell plate (pore size = 8 μm). Chemoattractant (medium with 10% FBS) was added to lower chamber. For invasion assays, Matrigel stock solution (Corning) was diluted with sterile ice-cold deionized water (1:4). Migrated cells were stained with crystal violet. For scratch assays, cells were seeded at 95% confluence on 6-well plates. Cell layers were scratched with P200 tips. Wound closure was determined by light microscopy.

### In vivo studies

All animal experiments were approved by Massachusetts General Hospital’s Institutional Animal Care and Use Committee (IACUC). 6–7-week-old female athymic nude mice were purchased from Charles River Laboratories. Animals were housed in a 12-hour light-dark cycle with access to food and water *ad lib*.

### Spontaneous models of BM: Intracardiac tumor injection

CALU6-FmC (NDUFA10•WT or •C253S) and NCIH661-FmC (NDUFA10•WT or •C253S) cells (*17*) were pelleted and resuspended in Hank’s Buffered Saline Solution (HBSS). Mice were anesthetized using 3% isoflurane, and a 50 µL cell suspension containing 1 × 10^7^ CALU6-FmC or 5 × 10^5^ NCIH661-FmC cells was injected into the left cardiac ventricle using a 29 g 1/2mm insulin syringe (n=9-10 animals per group). After 30 days, mice were imaged at the MGB Center for Systems Biology (CSB) Imaging Core in prone and then supine positions (5 minutes each). Briefly, animals were anesthetized using 3% isoflurane. D-luciferin solution was prepared in saline and administered to mice at 4.5 mg/kg via intraperitoneal injection. After 10 minutes, *in vivo* images were acquired using AMI HTX optical imaging platform (Spectral Instruments Imaging). Aura *in vivo* Imaging Software (Spectral Instruments Imaging) was used to define regions of interest (ROI) and quantify tumor burden (as total radiance above background signal, normalized on ROI area) for each animal. Immediately after *in vivo* imaging, animals were euthanized, and brains and lungs were collected. *Ex vivo* imaging was performed on these organs (3-5 minutes). *Ex vivo* images were blinded and reviewed by the MGH CSB Imaging Core to determine tumor positivity or negativity and assess BM.

### Orthotopic models of BM: intracranial tumor injection

NCIH661-FmC (NDUFA10•WT or •C253S) cells were pelleted and resuspended in HBSS. Then, 2 µL of cell suspension containing 2.5 × 10^5^ NCIH661-FmC cells were injected into the right brain hemisphere, respectively, to generate intracranial models of BM as previously described (*17*) (n=10 animals per group). Bioluminescence imaging (BLI) at the MGB Center for System Biology Imaging Core was used to monitor intracranial tumor growth at different time points.

### Analysis of cell lines

#### Analysis of cysteine chemical proteomics and unenriched proteomics

For **unenriched proteomics** (Supplementary Table S2), raw MS3 intensities were first normalized using the size factors method (*135*) after the addition of a pseudocount of 1. Log₂ fold changes (treatment/vehicle) were then calculated using the normalized intensities. For significance testing of protein expression changes, an empirical null distribution was estimated using vehicle controls by calculating the log₂ fold changes between vehicle pairs (H₂O vs. DMSO and DMSO vs. H₂O). Treatment-associated log₂ fold changes were compared against the two empirical null distributions using a two-sample t-test, and the more conservative (larger) p-value was retained. P-values were subsequently adjusted for multiple hypothesis testing using the Benjamini-Hochberg procedure.

For **cysteine chemical proteomics** (Supplementary Table S1), raw intensities were not normalized, and a pseudocount of 1 was added prior to calculating log₂ fold changes (treatment/vehicle). The lack of normalization distinction stems from the experimental design: in TMT-based protein-level proteomics, equal protein amounts are loaded across channels, and variation in total signal is assumed to arise primarily from technical error. In contrast, chemical proteomics involves an enrichment step (e.g., desthiobiotin-iodoacetamide-based enrichment), which disrupts the assumption of equal MS3 intensity across channels. To emphasize relative changes while reducing the influence of technical variability, log₂ fold changes were z-scored across TMT channels (individual samples). For significance testing of cysteine accessibility changes, the resulting accessibility scores for each cysteine were tested against a null hypothesis of zero mean using a t-test.

### Correcting for protein expression changes

Two perturbations (DOXY and KI696) were profiled following 48 hours of treatment, a duration deemed sufficient to induce substantial changes in protein abundance that could confound subsequent interpretation of cysteine accessibility measurements. For example, cysteines on GCLC appear to increase in accessibility following NRF2 activation; however, this effect can be explained by a substantial increase in total GCLC protein abundance, resulting in a larger pool of detectable cysteines independent of changes in post-translational regulation.

To account for protein expression changes, 19 of the 55 profiled cell lines underwent both protein-level and cysteine-level profiling (Supplementary Table S2). To eliminate proteins whose abundance changes could influence the accessibility readout of long-term treatments, which are more likely to cause protein expression changes, cysteines mapping to proteins with a significant change in abundance (adjusted p-value ≤ 0.05) or whose accessibility changes were significantly correlated with the corresponding protein abundance change (Pearson r ≥ 0.4 and p-value < 0.05) were flagged and removed from downstream analysis for DOXY and KI696 treatments.

### Cysteine enrichment analysis

#### Cysteine-level enrichment for known pathways

cysteine enrichment analyses (CSEA) were conducted using a custom MATLAB script implementing the unweighted GSEA scoring function as previously described (*135*). False discovery rates (FDR) were calculated using permutation-based null distributions, as described in the original GSEA publication (*135*). This strategy emphasizes rank-based enrichment rather than fold-change magnitude, which can be more variable in noisy datasets.

Cysteine sets were compiled (Supplementary Table S3) from KEGG (*136*), Reactome (*137*), and PantherDB (*138*). Previously curated cysteine sets from prior manuscript were also used (*53*). For the chemical proteomics enrichment analysis in the cell lines data, cysteines detected in at least 20 biological samples were ranked by absolute accessibility change weighted by the negative log10-transformed p-value, and enrichment was performed on the resulting ranked list. Essentially, this reflects the overall burden of protein-state changes within a given pathway or organelle.

### Determining protein state signatures

To construct **oxidative and reductive protein state signatures** (Supplementary Table S4), cysteines were first filtered to require detection in at least 20 H₂O₂-treated samples and 20 reductive-perturbation samples, yielding just under 10,000 cysteines. Significant cysteines were then selected using a Bonferroni-scale threshold of p-value < 1 × 10⁻□ and a median accessibility change ≥ 0.5 or ≤ −0.5 in at least one treatment condition. The oxidative signature was derived from H₂O₂ treatment, whereas the reductive signature was derived from KI696, DOXY, GSH, NAC, and Trolox treatments. Each signature consisted of an "upper" and a "lower" component, corresponding to cysteines that exhibited significantly increased or decreased accessibility, respectively. For long-term reductive perturbations, DOXY and KI696, cysteines mapping to proteins with significant protein abundance changes or protein-correlated accessibility changes were excluded before signature construction. To remove ambiguous cysteines, cysteines were excluded if they were assigned to both the upper and lower components within the same signature class, or if they were assigned to the same direction in both oxidative and reductive signatures.

To construct perturbation-specific significant cysteines were defined by a p-value < 1 × 10⁻O. For organelle-specific signatures, cysteines were assigned to organelles based on UniProt subcellular localization annotations with the above p-value thresholds. Condition-specific and organelle-specific signatures were then constructed using the same procedure described above.

### Analysis of human samples

#### Analysis of untargeted proteomics and cysteine chemical proteomics

For protein-level proteomics, raw MS3 intensities were first normalized using the size factors method (132), after which a pseudocount of 1 was added. No size factor normalization was performed for cysteines. Both measurements were normalized to their corresponding bridge channels in log₂ space to account for batch effects across TMT plexes. Following bridge normalization, each individual TMT channel was z-scored across all quantified proteins or cysteines to emphasize relative differences and reduce the influence of technical variation. Each tumor sample was profiled in triplicate, and replicate measurements were collapsed by taking the median, yielding a single protein-level and cysteine-level measurement per sample. To account for the contribution of protein abundance to cysteine accessibility measurements, a separate linear model was fit for each cysteine across all tumor samples using the abundance of its corresponding protein as the predictor and cysteine abundance as the response variable. The residuals from these models were retained and referred to as expression-corrected accessibility values.

#### Correcting for tumor purity

To prevent non-tumor cell-type contributions from confounding our results in low-purity samples, we identified cysteines whose accessibility varies across different contributions of normal cell types and excluded them from downstream analysis. Reasoning that the majority of non-tumor signal in primary tumors would arise from infiltrating immune and stromal cells, as described previously (*139*), and in addition from brain-derived cells in brain metastases, we derived protein expression signatures for these tissues. Candidate stromal and immune marker genes were obtained from CellMarker2.0 database (*140*). Immune signals are composed of markers of T cells, B cells, NK cells, plasma cells and other myeloid-derived cells. Stromal signals are composed of markers of endothelial cells, myofibroblasts and pericytes. Brain-specific markers were obtained from the Human Protein Atlas and defined as genes exhibiting at least five-fold higher expression in brain tissue relative to other tissues (*141*).

Marker genes whose corresponding proteins were detected in 100% of the samples samples were retained. To improve marker specificity, pairwise gene-gene correlations were calculated across all primary lung tumors, normal lung samples, and brain metastases. Genes that failed to positively correlate with other members of their respective marker set were excluded. The remaining marker genes were used to construct stromal, immune, and brain signatures by calculating the mean protein abundance of each marker set within each sample (fig. S2B-C).

To identify features potentially influenced by non-tumor cellular composition, Spearman correlations were calculated between each protein or cysteine and the stromal, immune, and brain signatures. Correlations were evaluated separately within lung specimens (normal and primary tumors) and brain metastases. P-values were adjusted using the Benjamini-Hochberg procedure. Proteins or cysteines exhibiting a significant association with stromal or immune signatures in either cohort (|ρ| ≥ 0.6 or adjusted p-value ≤ 0.05) or with the brain signature within brain metastases were flagged as potentially confounded by cellular composition and excluded from downstream analyses.

#### Determination of relative oxidation

To quantify protein ROS states in patient tumors, expression-corrected cysteine accessibility values were first normalized to the median of the normal lung samples for each cysteine. Single-sample cysteine set enrichment analysis (ssCSEA) was then performed independently for each tumor using the oxidative, reductive (union set of cysteines detected across all reductive conditions), condition-specific, and organelle-specific signatures described above (Supplementary Table S3).

Each signature consisted of paired increased-accessibility ("upper") and decreased-accessibility ("lower") cysteine sets. For each sample, ssCSEA generated a normalized enrichment score (NES) for both components. Oxidative and reductive signature scores were calculated by subtracting the NES of the decreased-accessibility component from the NES of the increased-accessibility component:

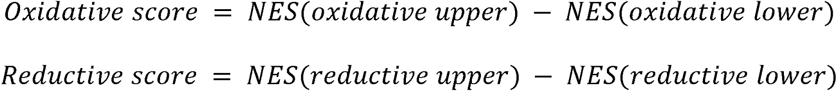

A relative oxidation score was then calculated as:

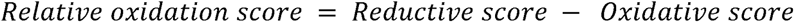

Positive relative oxidation scores indicate enrichment of cysteine accessibility patterns associated with oxidative perturbations, whereas negative scores indicate enrichment of cysteine accessibility patterns associated with reductive perturbations. The same framework was applied to condition-specific and organelle-specific signatures. To test for differences in ssCSEA-derived redox scores between non-metastasized primary tumors, metastasized primary tumors, and brain metastases, one-tailed Wilcoxon rank-sum tests were performed for both possible directions of change, and the smaller p-value was retained.

#### Fingerprint plots

To visualize the structure of cysteine reactivity across tumors, expression-corrected cysteine accessibility values were first normalized to the median of the normal lung samples for each cysteine. The resulting tumor-level profiles were z-scored (across tumors) and embedded into two dimensions using t-SNE. The resulting (X, Y) coordinates were transformed into concentric, sinusoidally distorted elliptical rings. This transformation involved centering the data, converting to polar coordinates, binning points into radial “layers” based on distance from the origin, and projecting them back into Cartesian space with sinusoidal radial perturbations and anisotropic scaling. The resulting embedding emphasizes radial structure while preserving local neighborhood relationships.

#### Determining response to ROS-lowering treatments

To quantify the responsiveness of each cell line to antioxidant treatment, cysteine accessibility changes were aggregated across replicates for each treatment by taking the median accessibility score for each cysteine. Reductive state signatures were then scored using ssCSEA as described above. For each cell line and treatment, a reductive response score was calculated as:

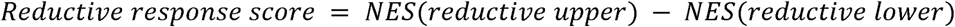

To obtain a treatment-independent measure of antioxidant responsiveness, reductive response scores were averaged across DOXY, GSH, KI696, NAC, and Trolox treatments. Cell lines with larger positive scores were considered more responsive to ROS lowering perturbations.

#### Analysis of genomic screens

The raw read matrices for genomic screens were corrected for library complexity using size factor normalization (*142*), after adding a pseudo-count of one. Scores were then calculated as the log₂ fold change of treatment relative to vehicle control. For the CRISPR screen, gene scores were filtered to retain only genes with two or more detected guides.

### Gene enrichment analysis

***Gene-level enrichment:*** Gene Set Enrichment Analysis (GSEA) for the CRISPR knockout screens was performed using the *fgsea* R package (*143*) on all MSigDB pathway gene sets (*22*).

### Identification of functional cysteines responsive to lower ROS

To identify cysteines associated with lower ROS sensitivity, cysteine accessibility changes from ROS-lowering perturbations (DOXY, GSH, KI696, NAC, and Trolox) were integrated with CRISPR knockout screening data. In this study we focused on mitochondrial genes given the strong overall enrichment of genes localizing to this organelle from CRISPR screens (Figure XXX). Mitochondrial oxidative phosphorylation genes were obtained from MitoCarta 3.0 (*144*) and cysteines were mapped to their corresponding genes. For each cysteine, the ROS-lowering perturbation producing the largest absolute accessibility change was identified after filtering for cysteine detection and excluding protein abundance-confounded measurements as described above. The accessibility change under this maximally responsive condition was then paired with the CRISPR knockout phenotype measured under the corresponding treatment condition. Cysteines were subsequently ranked based on the combination of reductive accessibility response and genetic dependency.

### Statistical analysis

Statistical analyses were performed with and Prism v10 (GraphPad). Error bars represent mean ± SD. Statistical comparisons were analyzed using two-tailed Student’s *t*-test or one-tailed Wilcoxon rank-sum test with P values indicated in figure legends and source data. The Fisher’s exact test was performed to the animal experiments. A False Discovery Rate (FDR) was calculated for Proteomics (using Proteome Discover, v2.5, Thermo Fisher Scientific) analysis to correct for multiple comparisons. Data were considered statistically different at * *P* < 0.05, ** *P* < 0.01, and *** *P* < 0.001.

